# A Small Multidrug Resistance Transporter in *Pseudomonas aeruginosa* Confers Substrate-Specific Resistance or Susceptibility

**DOI:** 10.1101/2023.09.28.560013

**Authors:** Andrea K. Wegrzynowicz, William J. Heelan, Sydnye P. Demas, Maxwell S. McLean, Jason M. Peters, Katherine A. Henzler-Wildman

## Abstract

Small Multidrug Resistance (SMR) transporters are key players in the defense of multidrug-resistant pathogens to toxins and other homeostasis-perturbing compounds. However, recent evidence demonstrates that EmrE, an SMR from *Escherichia coli* and a model for understanding transport, can also induce susceptibility to some compounds by drug-gated proton leak. This runs down the ΔpH component of the Proton Motive Force (PMF), reducing viability of the affected bacteria. Proton leak may provide an unexplored drug target distinct from the targets of most known antibiotics. Activating proton leak requires an SMR to be merely present, rather than be the primary resistance mechanism, and dissipates the energy source for many other efflux pumps. PAsmr, an EmrE homolog from *P. aeruginosa*, transports many EmrE substrates in cells and purified systems. We hypothesized that PAsmr, like EmrE, may confer susceptibility to some compounds via drug-gated proton leak. Growth assays of *E. coli* expressing PAsmr displayed substrate-dependent resistance and susceptibility phenotypes, and *in vitro* solid-supported membrane electrophysiology experiments revealed that PAsmr performs both antiport and substrate-gated proton uniport, demonstrating the same functional promiscuity observed in EmrE. Growth assays of *P. aeruginosa* strain PA14 demonstrated that PAsmr contributes resistance to some antimicrobial compounds, but no growth defect is observed with susceptibility substrates, suggesting *P. aeruginosa* can compensate for the proton leak occurring through PAsmr. These phenotypic differences between *P. aeruginosa* and *E. coli* advance our understanding of underlying resistance mechanisms in *P. aeruginosa* and prompt further investigation into the role that SMRs play in antibiotic resistance in pathogens.

**IMPORTANCE:** Small multidrug resistance transporters are a class of efflux pumps found in many pathogens, but whose contributions to antibiotic resistance are not fully understood. We hypothesize that these transporters may confer not only resistance, but also susceptibility, by dissipating the proton-motive force. This means to use an SMR transporter as a target, it merely needs to be present (as opposed to being the primary resistance mechanism). Here, we test this hypothesis with an SMR transporter found in *Pseudomonas aeruginosa* and find that it can perform both antiport (conferring resistance) and substrate-gated proton leak. Proton leak is detrimental to growth in *E. coli* but not *P. aeruginosa*, suggesting that *P. aeruginosa* responds differently to or can altogether prevent ΔpH dissipation.

## INTRODUCTION

Multidrug-resistant *Pseudomonas aeruginosa* is a threat to public health, causing over 30,000 hospital-acquired infections per year in the United States and an estimated 85,000 deaths worldwide in 2019 (1, 2). Some strains appear resistant to all known antibiotics (3). Due to the high degree of antimicrobial resistance and the limited availability of effective therapeutic treatment options, *P. aeruginosa* is a compelling and clinically relevant system to investigate mechanisms of drug resistance. One such mechanism is secondary active transport of toxic compounds by efflux pumps. The Small Multidrug Resistance (SMR) transporters are a family of efflux pumps that have been identified in many high priority pathogens, but their contributions to antibiotic resistance are minimally understood (4–11). The SMR family of transporters may be broadly divided into four subfamilies, with the QAC subfamily representing the most promiscuous transporters, named for their ability to confer resistance to <u>q</u>uaternary <u>a</u>mmonium <u>c</u>ompounds (11).

Historically, SMR transporters were understood as antiporters, tightly coupling proton import to drug export (12, 13). However, *in vitro* biophysical investigation of the model QAC SMR transporter EmrE has demonstrated that EmrE is also theoretically capable of symport and drug or proton uniport, with transporter behavior determined by substrate identity and kinetic parameters (14–17). This differential transport behavior, termed the “free-exchange model,” will ultimately lead to the transporter conferring susceptibility to some chemicals instead of resistance (see Figure 3A).

Substrate-gated proton uniport has been experimentally demonstrated for EmrE in *E. coli*, leading to disruption of the ΔpH component of the proton-motive force (PMF), impacting cellular ATP production and other PMF-dependent processes (15). PMF disruption has been theorized to be a potential antibiotic and/or adjuvant strategy (18–22). Should other SMR transporters or even other families of efflux pumps confer opposing biological outcomes in a substrate-dependent manner, this may broaden our understanding of how to target pathogens, inducing susceptibility instead of merely disabling specific bacterial resistance processes.

PAsmr, also known as EmrE_Pae_, is an SMR transporter expressed by *P. aeruginosa,* originally identified on the basis of sequence homology to EmrE and conferring resistance to ethidium bromide, methyl viologen, and other quaternary ammonium compounds (4, 7, 23, 24). *smrPA* (PAO1_4990, PA14_65990), the gene expressing PAsmr, has been identified in clinically isolated strains of multidrug resistant *P. aeruginosa* (25). PAsmr has also been investigated as a potential target and model for efflux pump inhibiting peptides (23). We previously carried out a chemical sensitivity screen expressing PAsmr in *E. coli*, demonstrating that PAsmr displays functional promiscuity, the ability to confer either resistance or susceptibility depending on which substrate is present (24). This is similar to the behavior observed for EmrE (15).

Here, we further explore the ability of PAsmr to confer either resistance or susceptibility depending on substrate, and hypothesize that these opposing biological outcomes are explained by different transport modes, based on our prior observations of EmrE (14, 15, 17). We use heterologous expression in *E. coli* to assess PAsmr activity and find that it can confer either resistance or susceptibility to different substrates. This confirms the phenotypes previously identified in an unbiased screen (24). We use *in vitro* assays to demonstrate that the resistance phenotypes are due to proton-coupled antiport, while substrate-triggered proton uniport is responsible for susceptibility phenotypes. Finally, we use a *P. aeruginosa* strain lacking PAsmr to look for phenotypic differences in the presence of both known and novel SMR substrates (Figure 1) in the native organism, as well as how this phenotype corresponds to those observed upon heterologous expression in *E. coli*. The results highlight differences in the impact of proton-coupled transport and proton leak in *E. coli* and *P. aeruginosa*.

**Figure 1.**
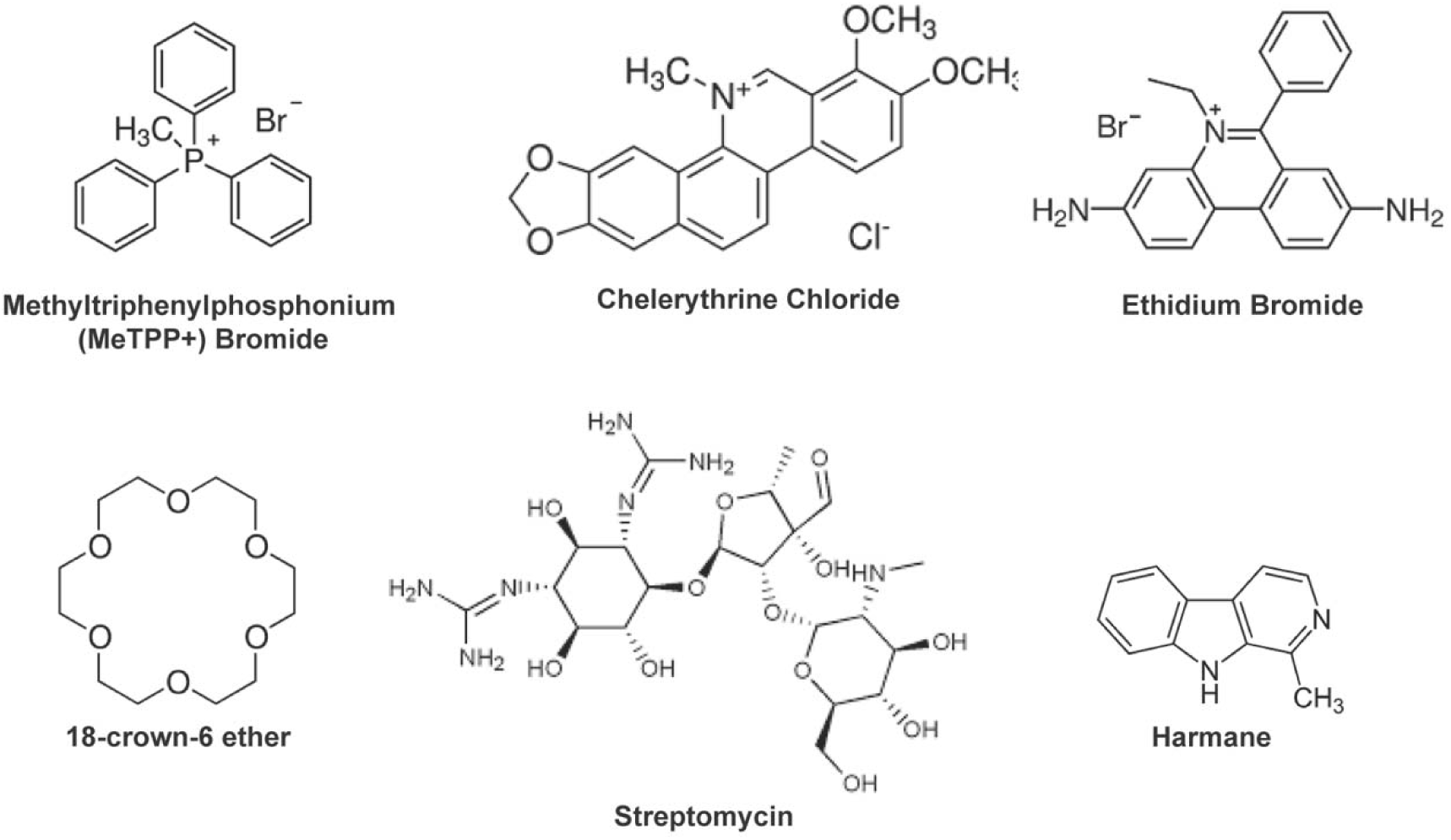
Known and novel PAsmr substrates. Compounds used in this study.

## MATERIALS AND METHODS

### Growth assays in E. coli

MG1655 Δ*emrE E. coli* cells were transformed with either WT or non-functional (E14Q) PAsmr (PAO1-PA4990) cloned into the pWB plasmid, a low copy, leaky-expression vector (24). Cells were grown in Nutrient Broth (Difco 234000),100 µg/mL carbenicillin, from a single colony to an OD of 0.2 at 37°C. Cultures were diluted to a final OD of 0.01 in microplates (Corning, REF: 351172) containing select concentration values of tested compounds. The plates were incubated in a microplate reader (BMG-Labtech or TECAN) at 37 °C. OD_600_ (or OD_700_ for ethidium) was measured every 5 minutes for 18 h. Experiments were performed in biological triplicate.

### ATP Quantitation in *E. coli*

ATP quantitation was carried out using the commercially available BacTiter-Glo^TM^ Assay from Promega. Cells were grown as described in the growth assays until 18 h (or 6 h when specified), and then transferred to a black 96-well plate and mixed with the BacTiter-Glo^TM^ Reagent. Plates were shaken for 5 minutes at RT and luminescence detected with a TECAN microplate reader.

Experiments were performed in biological triplicate. Statistical significance was determined by Student’s t-test and based on p < 0.05.

### PAsmr Expression and Purification

PAsmr was expressed as a maltose binding protein (MBP) fusion construct, using pet28-TEV-MBP, a gift from Zita Balklava & Thomas Wassmer (Figure S5A) (26). Following TEV protease cleavage of the N-terminal MBP, the N-terminal 6xHis-tag may be used to purify PaSMR via affinity chromatography, and then removed with thrombin leaving an extra N-terminal GSHGS. BL21(DE3) Gold cells transformed with pet28-MBP-PaSMR were grown in M9 minimal media (27), with protein expression induced by 0.33 mM IPTG at an OD_600_ of 0.9-1 at 17°C. Cells were harvested after 17 hours and cell pellets were resuspended in lysis buffer (250 mM sucrose, 100mM NaCl, 2.5 mM MgSO_4_, 20mM tris pH 7.5, 5mM β-mercaptoethanol, 1 mg/mL lysozyme, DNAse, 1 μg/mL pepstatin, 10 μM leupeptin, and 100 μM PMSF), and lysed by sonication. 0.25 mg/mL TEV protease was added, and following incubation for 16-48 h at RT or 48-72 h at 4°C, the membrane fraction was separated by a high-speed spin (30,000 × *g* for 1), resuspended in the same buffer, and solubilized with 40 mM DM (decylmaltoside, Anatrace) at RT for 2 hours. After a second high-speed spin, 6x-His tagged PAsmr was purified via NiNTA affinity chromatography, thrombin cleavage, and size-exclusion chromatography as previously described for EmrE (Figure S5B, C, E) (27). When more complete removal of MBP was required for downstream experiments (e.g. SSME), 50-100 mM maltose was included in all steps of purification through NiNTA column elution and an additional desalting step was added to the end of the purification. Identity of PAsmr as the purified species was confirmed by whole-protein mass spectrometry analysis (Figure S5D).

### Solid-supported membrane electrophysiology

Purified WT PAsmr was reconstituted into 1-palmitoyl-2-oleoyl-sn-glycero-3-phosphocholine (POPC) liposomes at a lipid-to-protein molar ratio of 250:1 in pH 7 assay buffer (100 mM Potassium Phosphate, 100 mM Sodium Acetate, 200 mM NaCl, 4 mM MgCl_2_). Liposome preparation was performed as described in (15). All SSME data were acquired using a Nanion SURFE2R N1 instrument as described in (28). Liposome aliquots were thawed, diluted 2-fold with assay buffer, and briefly sonicated. 10 μL of liposomes were used to prepare 3 mm sensors as previously described (15). Before experiments, sensor capacitance and conductance values were obtained to ensure sensor quality. All experiments used assay buffer with internal pH values of 7.0 and external pH values of 6.7. Drug concentrations used are in Table S1. Sensors were rinsed with at least 500 μL of internal buffer before each measurement to set the internal buffer, pH, and drug concentrations as described in (28). Data acquisition occurred in three stages. First, sensors were perfused with an internal buffer, then transport was initiated by perfusion of the external buffer, and finally, perfusion of the internal buffer re-equilibrated the sensors. Signals were obtained by integrating the current during perfusion of the external buffer, with the final 100 ms of the initial internal buffer perfusion used as the baseline. Reported data are average values of at least three sensors, with error bars representing the standard deviation. Statistical significance was determined by Student’s t-test and based on p < 0.05.

### 1D Ligand-Observed NMR

PAsmr was purified as described above. Binding of MeTPP^+^ and streptomycin was assessed by ^1^H NMR. Samples contained 500 µM MeTPP^+^ or streptomycin with or without 25 µM PAsmr in DHPC/DMPC bicelles, in a buffer of 20 mM NaCl, 20 mM potassium phosphate, 8% D_2_O, and 0.3 mM sodium trimethylsilylpropanesulfonate (DSS) at pH 7. Samples containing protein also contained 0.05% sodium azide and 2 mM tris (2-carboxyethyl)phosphine (TCEP). All spectra were acquired on a Bruker Avance III spectrometer operating at 600 MHz. 1D spectra were acquired using the Bruker pulse program zgesgp. WaterLOGSY experiments were recorded at specified temperatures using the Bruker pulse program ephogsygpno.2 with a mixing time of 1.5 s and 2 s recycle delay. All spectra were processed and visualized with Mnova.

### Proton Leak

PAsmr was reconstituted into 3:1 POPC:POPG liposomes at a lipid-to-protein molar ratio of 400:1. Proton leak was monitored by diluting PaSMR liposomes in inside buffer (50 mM sodium phosphate, 100 mM KCl, pH 7) into weakly buffered outside buffer (75 μM Phenol Red, 99 mM NaCl, 1 mM KCl, pH 7) to 0.8 µM final PaSMR concentration. Valinomycin diluted in inside buffer was added to final concentration of 1 µg/mL to create a negative inside membrane potential. FCCP diluted in inside buffer was added to final concentration of 1 µg/mL as a positive control. At the end of the assay, HCl (50 nM final) was added to return to resting potential.

### smrPA (PA14_65990*) Deletion in* P. aeruginosa *PA14*

*smrPA* was deleted in PA14 using *sacB* counterselection (29, 30). Briefly, a plasmid vector (pJMP3262) containing an origin of replication that replicates in *E. coli* but not *P. aeruginosa*, a gentamicin resistance cassette, and the *sacB* gene was used as the backbone. The regions 1000bp upstream and downstream of *smrPA* were cloned into the plasmid vector via Gibson Assembly.

Following extraction of genomic DNA from PA14, the regions 1000bp upstream and downstream of *smrPA* were amplified via PCR using primers oJMP 1832 and 1833 for the upstream region and oJMP1834 and 1835 for the downstream region (Table 1). The resulting products were gel purified and cloned into pJMP 3262 via Gibson Assembly. The plasmid was then transformed into *E. coli* (sJMP 2630). The plasmid sequence was confirmed by Sanger sequencing using primers oJMP 1339,1340,1836, and 1837 (Table 1). The plasmid was then transformed into a conjugative *E. coli* strain (sJMP3257).

**Table 1.**
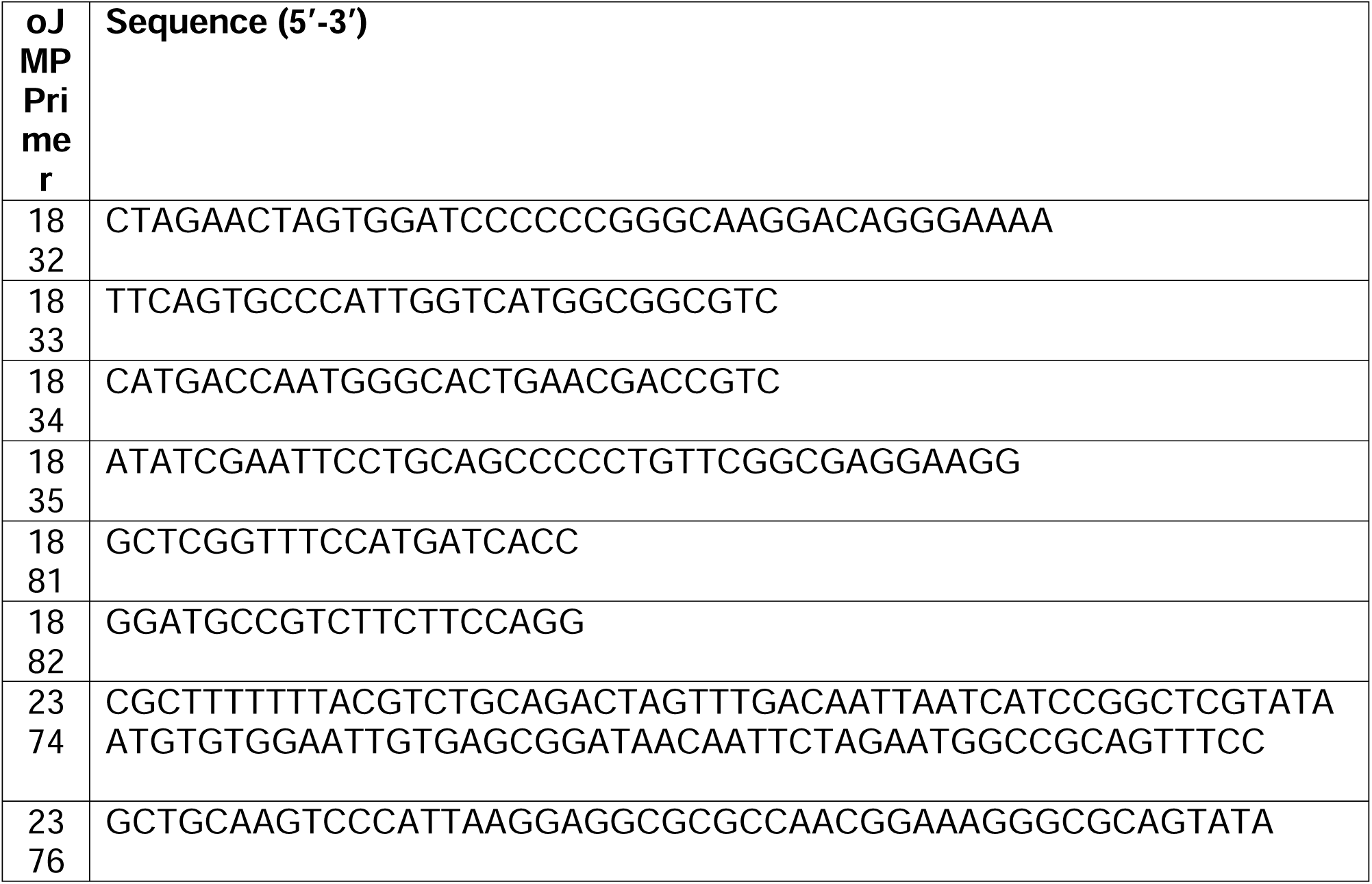
Primers used in PAsmr deletion and complementation.

The newly constructed plasmid donor (sJMP3257) and recipient (WT PA14 sJMP744) were grown overnight on plates and resuspended in 1mL LB to OD_600_ ≍ 3. Three tubes were prepared, one for mating containing 800 µL LB, 100 µL of the recipient, and 100 µL of the donor strain, and two controls (900 µL LB and 100 µL of either donor or recipient strains alone). Cells were pelleted at 4000 x *g* for 3 minutes. Media was removed and the pellet was resuspended in 25 µL of LB and spotted on a filter on a LB + diaminopimelic acid (DAP) plate. Following incubation for 3.5 h at 37°C, the filter was placed in a 1.5 mL Eppendorf tube with 200 µL LB and vortexed for 30 s. The cells were further diluted in LB at 1:10, 1:100, 1:1000, 1:10000, and 1:100000. 90 µL of each dilution was plated onto LB + gent (30 µg/mL) plates to select for *P. aeruginosa* transconjugants containing the plasmid integrated into the genome at the *smrPA* (PA14_65990) locus by single crossover. Donor and recipient controls were also plated on LB + gent (30 µg/mL) plates.

Several isolated colonies from the LB + gent (30 µg/mL) plates were grown in 10 mL LB overnight. The resulting overnight cultures were plated on LB + sucrose 15% plates and incubated for 48-72 h at room temperature (RT). Incubation at RT enhanced recovery of correct, double-crossover clones. To confirm the selection of Δ*smrPA*, 40 isolated colonies from the LB +sucrose 15% plates were patched on LB + gent (30 µg/mL) + sucrose 15%, LB + gent (30 µg/mL), and LB plates. Any colonies growing on the plates containing gent (30 µg/mL) were discarded. To confirm the deletion of *PAsmr,* the gDNA was extracted from a few samples, and the primers oJMP1881 and oJMP 1882 were used to amplify the region including *smrPA.* Gel electrophoresis of the PCR product was used to confirm the deletion of the 333 bp gene product (Figure 6A).

### smrPA (PA14_65990*) Complementation in* P. aeruginosa *PA14*

An isopropylthiogalactoside (IPTG) inducible promoter, Ptrc, was chromosomally inserted upstream of *smrPA* in the PA14 Δ*smrPA* strain background to complement its function. The plasmid expression vector, pJMP 3389, which contains an apramycin (*P. aeruginosa* selection) and ampicillin resistance (*E. coli* selection) cassette was used to create the *smrPA* complemented strain. The PCR primers oJMP 2374 (containing P*_trc_*) and oJMP 2376 (Table 1) were used to amplify PA14 genomic DNA and create the Ptrc-*smrPA*^+^ insertion. Following a gel purification of the PCR product and restriction digest of the mini-Tn*7* plasmid pJMP3389, the Ptrc*-smrPA*^+^ insert was cloned into the digested plasmid vector via Gibson assembly.

The newly assembled plasmid vector was then transformed into *E. coli* (sJMP 2630) on LB + ampicillin (100 μg/mL) selective plates, and the resulting plasmid sequence was confirmed by whole plasmid sequencing (Nanopore sequencing from Plasmidsaurus). The plasmid was then transformed into a conjugative *E. coli* strain (sJMP3257). In a tri-parental mating to integrate Tn*7* containing the P*_trc_*-*smrPA*^+^ cassette into the *P. aeruginosa* Δ*smrPA* genome, newly constructed plasmid donor (sJMP3257), recipient (PA14 ΔsmrPA), and transposase donor (sJMP 2953) were grown on agar plates overnight and resuspended in 1mL of LB to OD_600_ ≈15, OD_600_ ≈5.5, and OD_600_ ≈12, respectively. Four tubes were prepared, one for mating containing 700uL LB, 100uL of the recipient, 100uL of the transposase donor, and 100uL of the donor strain, and three controls (no recipient, no transposase donor, and no transposon donor controls). The cells were pelleted, and the supernatant was removed. The resulting cell pellet was resuspended in 25uL of LB and spotted on a filter on a LB + diaminopimelic acid (DAP) plate. Following incubation for 3.5 hours, the filter was placed in a 1.5mL Eppendorf tube with 200uL LB and vortexed. The cells were further diluted in LB and plated on LB + Apramycin (50 μg/mL) selective plates.

### *Growth Assays in* Pseudomonas aeruginosa

WT PA14 and PA14 Δ*smrPA* were grown in Nutrient Broth (Difco 234000) from single colonies to an OD of 1 at 37 °C. Cultures were diluted to a final OD of 0.01 in microplates containing select concentration values of tested compounds. Plates were incubated in a microplate reader (TECAN) at 37 °C. OD_600_ (or OD_700_ for ethidium) were measured every 15 minutes for 18 h. Experiments were performed in at least biological triplicate. Data were analyzed by normalizing WT growth and standard deviation by growth of the knock-out strain.

## RESULTS

### E.coli expressing PAsmr display substrate dependent growth effects

Previously, we performed an unbiased screen to identify potential substrates of PAsmr using heterologous expression of the transporter in *Escherichia coli* (24). The screen identified, and subsequent experiments confirmed, several novel substrates including the small molecules harmane and 18-crown-6 ether. The screen also identified the aminoglycoside streptomycin and the antibiotic chelerythrine chloride as potential substrates of PAsmr. To further assess the opposing biological outcomes seen in the prior study, Δ*emre E. coli* expressing WT PAsmr were grown in Nutrient Broth (NB), a low-ionic-strength medium. As a control, we used *E. coli* expressing E14Q PAsmr. E14Q is a known transport-dead variant of EmrE (31), and E14Q mutation has been shown to abolish the ability of PAsmr to confer resistance to toxic compounds when heterologously expressed in *E. coli* (24). Therefore, the E14Q variant is a valuable negative control for phenotyping *E. coli* strains with heterologous expression of SMR transporters.

Expression of WT PAsmr conferred resistance to ethidium bromide (Figure 2A, S1A, B); methyltriphenylphosphonimum bromide (MeTPP^+^), which is closely related to known substrate TPP^+^ (Figure 2B, S1C, D) (4); and chelerythrine chloride (Figure 2C, S1 E,F). PAsmr expressed in *E. coli* conferred susceptibility to harmane (Figure 2D, S2A, B), 18-crown-6 ether, (Figure 2E, S2C, D), and streptomycin (Figure 2F, S2E, F). Several other putative hits identified in the prior screen showed no growth difference for *E. coli* expressing WT or E14Q PAsmr (Figure S3), demonstrating that not all hits identified based on NADH production show a significant growth phenotype. Compared to previously tested growth in higher ionic strength media (24), growth in nutrient broth also resulted in earlier and more pronounced phenotypic differences between WT and E14Q PAsmr (Figure 2, S1, S2, S4). PAsmr phenotypes for harmane and 18-crown-6 ether in low-ionic strength media are consistent but more pronounced than those in Muller-Hinton Broth (24), suggesting that decreased ability to compensate for ΔpH dissipation may occur in low-ionic strength media. Overall, for compounds tested previously in growth assays, phenotypic results were consistent with growth in *E. coli* in higher ionic strength media (24), and confirmed that PAsmr confers resistance to the known EmrE substrate, MeTPP^+^. The susceptibility to streptomycin in *E. coli* shown here is the first demonstration of PAsmr conferring susceptibility to an aminoglycoside antibiotic

**Figure 2.**
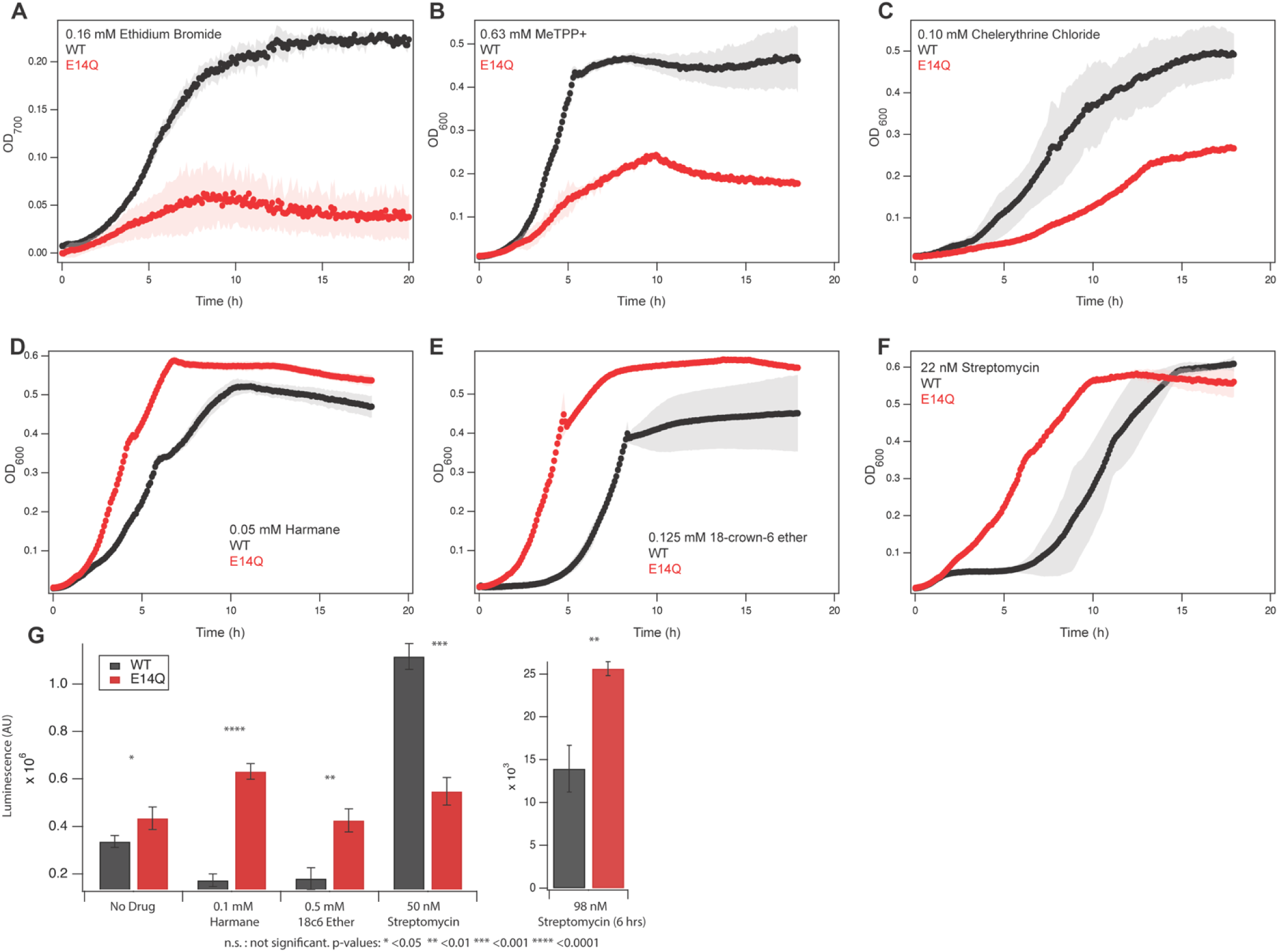
PAsmr expression in *E. coli* shows a substrate-dependent phenotype. A-C) *E. coli* expressing WT PAsmr shows improved growth relative to the E14Q variant in the presence of ethidium, MeTPP^+^, and chelerythrine, suggesting that PAsmr confers resistance to these compounds. D-E) E14Q-PAsmr-*E. coli* shows improved growth relative to WT PAsmr in the presence of harmane, 18-crown-6 ether, and streptomycin, suggesting that functional PAsmr confers susceptibility to these compounds in *E. coli*. G) ATP measurements of WT and E14Q PAsmr with harmane, 18-crown-6 ether, and streptomycin at 18 hours, and streptomycin at 6 hours. p-values were determined by Student’s t-test.

### ATP changes due to susceptibility are consistent with proton leak

While the functional promiscuity of PAsmr has been observed once previously (24), the molecular basis of susceptibility and the mechanism by which expression of functional PAsmr activity impairs growth of *E. coli* has not been established. We hypothesized that this susceptibility is due to substrate-grated proton leak through PAsmr, and this hypothesis is supported by the observation that some phenotypes are enhanced by a low ionic-strength medium. To further test this hypothesis, we also measured ATP levels of *E. coli* expressing PAsmr or non-functional E14Q PAsmr in the presence of the susceptibility substrates. If the PMF is dissipated due to proton leak, this should result in decreased levels of ATP production and increased levels of ATP consumption to restore PMF homeostasis (32).

ATP levels of *E. coli* expressing WT PAsmr and E14Q PAsmr were measured after 18 hours of growth. With no drug, E14Q PAsmr had slightly increased ATP levels, which may be explained by slight proton leak through PAsmr, or some basal transport by PAsmr that is detrimental to cell growth (Figure 2G). While the native role of Qac SMR transporters is not fully known, tolerance to pH and osmotic stress have been suggested as native functions, and may explain this slight growth difference (33, 34). Addition of harmane and 18-crown-6 ether resulted in significantly decreased ATP compared to the control, consistent with the proton leak hypothesis and the observed growth phenotypes (Figure 2G). In the presence of streptomycin, ATP levels of cells expressing WT PAsmr were increased, even though carrying capacity at endpoint was only slightly different from the transport-dead control (Figure 2F). To understand this result further, we measured ATP in the presence of streptomycin after 6 hours of growth, when the greatest difference in the growth phenotype was observed. After 6 hours, cells expressing E14Q PAsmr had significantly greater ATP levels than cells expressing WT PAsmr, consistent with the susceptibility phenotype.

Aminoglycosides have a complex and not fully understood mechanism of action. Inhibition of translation increases ATP levels by reducing ATP used for protein synthesis with time-dependent changes in ATP levels noted for *E. coli* exposed to aminoglycosides (35). In addition, changes in transmembrane voltage are implicated in aminoglycoside uptake and/or bactericidal activity (36, 37), adding complexity to how PAsmr-mediated proton leak and streptomycin activity may combine to influence the proton motive force and ATP levels.

### PAsmr substrate phenotype in E. coli corresponds to transport mode

Next, we investigated how PAsmr can confer opposing phenotypes depending upon substrate. We hypothesized that PAsmr may confer resistance via antiport but susceptibility via drug-gated proton uniport based on our prior studies of EmrE. To test if PAsmr performs only antiport or may perform drug-gated proton leak or other modes of transport (Figure 3A), we tested four substrates in a purified system using solid-supported membrane electrophysiology (SSME) (28). PAsmr was purified (Figure S5) and reconstituted into liposomes, which were adsorbed onto a gold electrode sensor. We note that PAsmr is an antiparallel homodimer with one subunit inserted in each orientation to form the functional dimer, removing any concern about the orientation of the protein when reconstituted into the liposome. In SSME, signal is detected as net charge moves in or out of the liposomes when different gradient conditions are applied. A two-fold outward pH gradient is established across the liposome in each assay, and the drug gradient is varied: Equal drug on both sides, greater drug concentration outside, or greater drug concentration inside (Figure 3B). Net charge transport in response to establishment of these gradients distinguishes between antiport, symport, and drug or proton uniport (Figure 3C).

**Figure 3.**
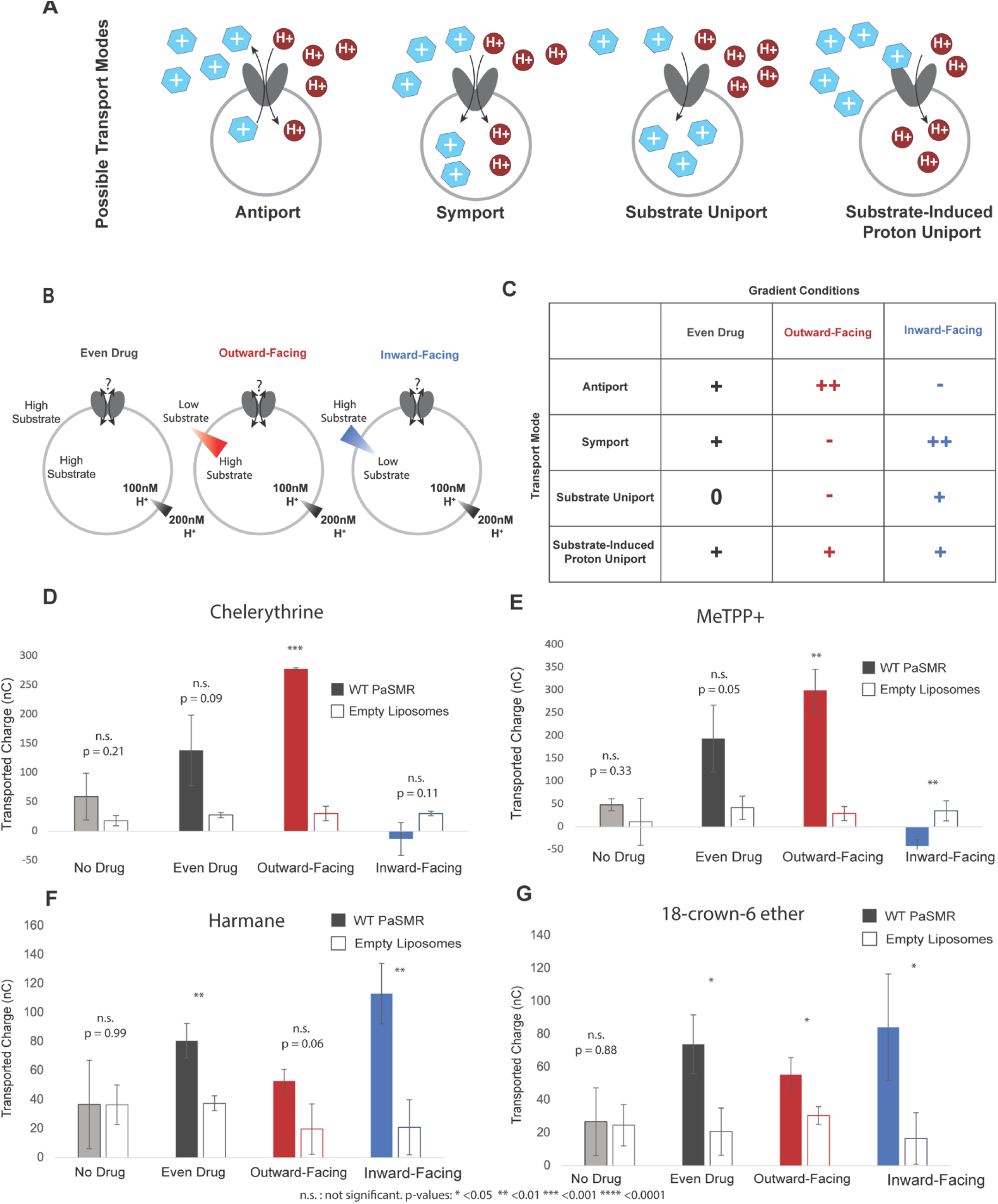
PAsmr performs both antiport and proton uniport depending on substrate. A) Based on the free-exchange model of EmrE, four transport modes are theoretically possible for PAsmr (16) B) Solid-supported membrane electrophysiology was performed using three different drug gradient conditions and a consistent 2-fold proton gradient. C) Resulting ion transport for each condition allows discrimination between four different transport modes. D) PAsmr transports Chelerythrine and E) MeTPP^+^ via antiport. F) Harmane and G) 18-crown-6 ether trigger increased proton transport regardless of drug gradient. p-values were calculated by Student’s t-test for each comparison between PAsmr and empty liposome, and are provided when below 0.15. Raw and integrated traces are available in Fig. S5 and S6.

In all cases, empty liposomes were used as a control, and background signal (2-fold proton gradient and no drug present) was not significantly different in PAsmr or empty liposomes, indicating that there is minimal proton leak through the liposomes and minimal proton leak through PAsmr proteoliposomes in the absence of drug. For all compounds, addition of drug in equal concentration inside and outside of the liposome resulted in a significant increase in transport compared to empty liposomes (Figure 3D-G, S6, S7). For MeTPP^+^ and chelerythrine chloride, addition of an outwardly directed 32x drug gradient resulted in the largest net charge transport (Figure 3C, S6D,E), and an inwardly directed 32x gradient resulted in reversal of charge, a hallmark of coupled antiport (28) (Figure 3D, E, S6D,F). This confirms that PAsmr performs proton-coupled antiport of chelerythrine and MeTPP^+^.

Harmane and 18-crown-6 ether, however, induced net positive transport under all conditions and does not reverse under any drug gradient conditions (Figure 3F, G, S7). This is consistent with harmane and 18-crown-6 ether not being transported, but instead triggering proton uniport down the 2-fold proton gradient (Figure 3C, S7). For the SSME experiments, we use substrate concentrations in a similar range to the concentration for which we observe phenotypes in bacteria. Unfortunately, in the case of streptomycin this concentration range is below the threshold where we can detect signal with SSME. We therefore used NMR to confirm that streptomycin binds to PAsmr.

### Ligand-detected NMR confirms streptomycin interaction

WaterLOGSY is a highly sensitive one-dimensional NMR technique that detects substrate binding under conditions where dissociation is fast (typically µM-mM affinity). 1H spectra are collected and free substrate will show positive signal while substrate or substrate regions bound to protein will show a negative signal (38). To confirm the WaterLOGSY effect with PAsmr, we first tested binding of MeTPP^+^ at 45 °C and found that MeTPP^+^ alone showed positive WaterLOGSY signal but upon addition of PAsmr and lipids, MeTPP^+^ and lipids both showed negative signal consistent with binding to PAsmr (Figure S8). Therefore, we used WaterLOGSY to confirm that streptomycin was interacting specifically with PAsmr. At 20 °C, the streptomycin signal in the WaterLOGSY experiment is positive when mixed with lipids alone. Upon addition of PAsmr, several streptomycin peaks became negative, consistent with specific moieties of streptomycin interacting with PAsmr (Figure 4). At lower temperatures, the full streptomycin spectrum became negative (Figure S9). The different temperatures used for MeTPP^+^ and streptomycin WaterLOGSY experiments reflect the difference in affinity of these two ligands and the requirement to have an off-rate on a suitable timescale for WaterLOGSY detection. This NMR data confirms that PAsmr interacts specifically with streptomycin.

**Figure 4.**
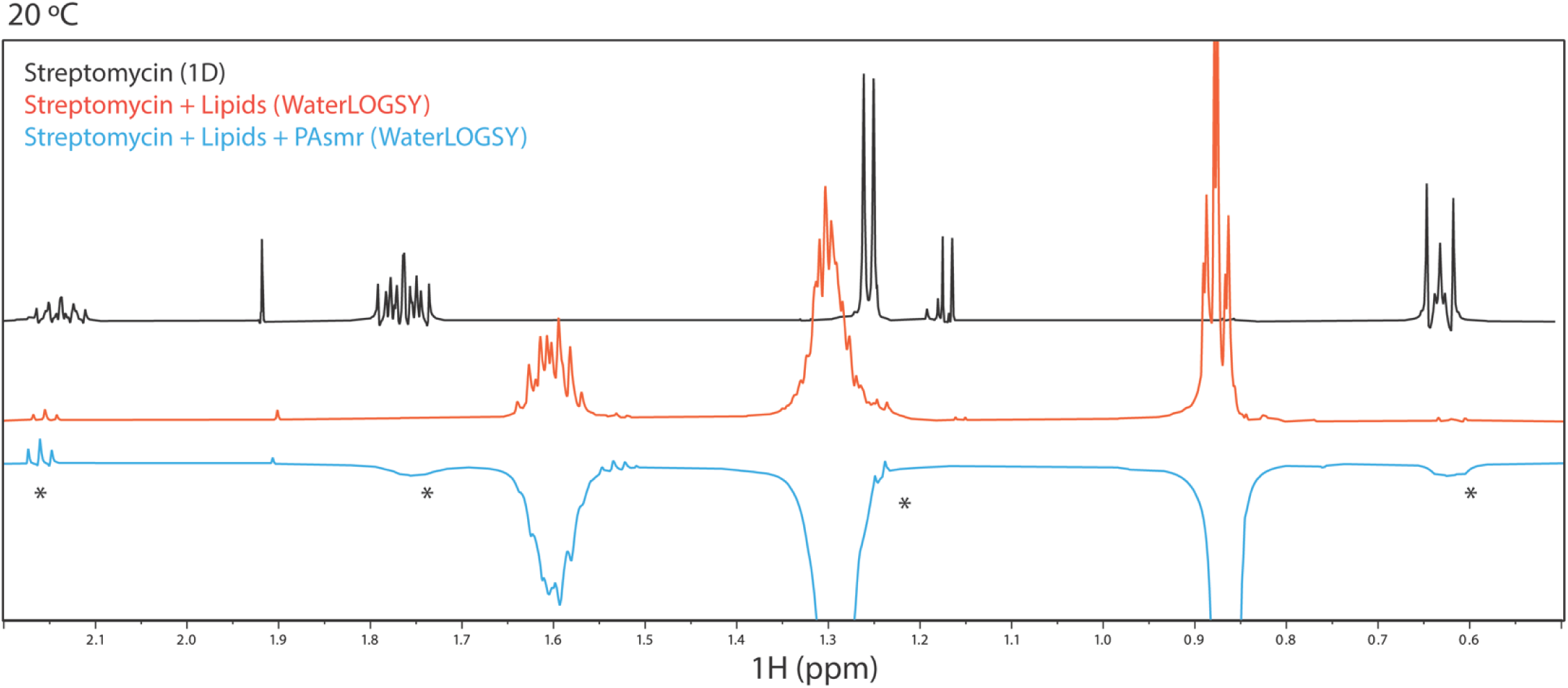
Streptomycin has selective interactions with PAsmr. Streptomycin has characteristic chemical shifts (black), and a positive WaterLOGSY signal with lipid alone (orange). Addition of PAsmr results in a negative WaterLOGSY signal at some but not all characteristic shifts (asterisk), confirming specific interaction of PAsmr and streptomycin. Additional spectra are shown in Fig. S8, S9, and available in the deposited data.

### Substrate is required to trigger proton leak

There was no evidence of proton leak in PAsmr liposomes with a proton gradient but no drug present (Figure 3, S6, S7); however, to confirm that the proton uniport observed in the SSME experiments was induced by drug and not simply due to potential inherent proton leak through PAsmr, we performed an additional voltage-driven proton leak assay as previously performed for EmrE (17). PAsmr was reconstituted in tight 3:1 POPC:POPG liposomes with a high internal potassium concentration at pH 7. These liposomes were then resuspended in a weakly buffered solution with a low potassium concentration and phenol red, which has pH-sensitive absorbance at 559 nm (Figure 5A). Addition of valinomycin creates a negative inside membrane potential as potassium moves down its concentration gradient. If PAsmr leaks protons, this negative inside membrane potential would drive proton leak through this pathway, causing the external pH to rise (lower proton concentration) leading to a change in Abs_559_ of phenol red. This did not occur (Figure 5B). The external pH shifted only after addition of a protonophore (FCCP) that allows direct proton transport across the membrane. These data indicate that PAsmr does not leak protons under a voltage gradient and substrate is required to trigger proton uniport or leak (Figure 5B, S10).

**Figure 5.**
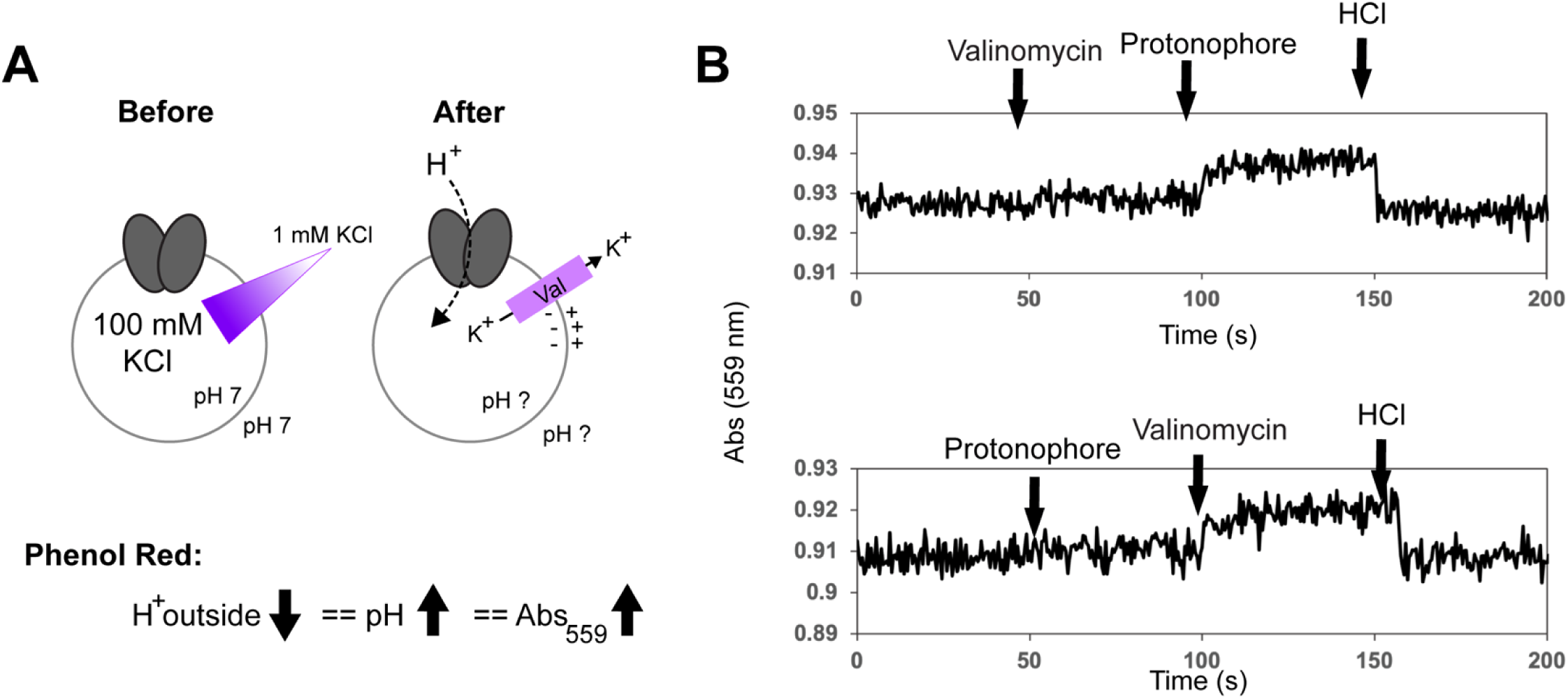
**PAsmr does not leak proton at neutral pH**. A) PAsmr was reconstituted into liposomes and resuspended with a 100-fold potassium gradient in neutral pH and phenol red. Addition of valinomycin dissipates the potassium gradient and establishes a negative-inside membrane potential, favoring proton transport into the liposome if it is possible. B) Phenol red absorbance at 559 nm reveals no change upon valinomycin addition (top), indicating that PAsmr does not leak proton under these conditions. Protonophore addition increases phenol red absorbance and HCl addition decreases absorbance, confirming establishment of a membrane potential. Protonophore addition first also does not result in an increase of absorbance, indicating that both membrane potential and protonophore are necessary for proton transport through liposomes (bottom). Representative traces are shown, and additional replicates are provided in Fig. S10.

### smrPA deletion phenotype in P. aeruginosa PA14 is substrate-dependent

Finally, to test the function of PAsmr and impact of proton leak in the native organism, we created a deletion strain in PA14, a clinically derived virulent strain of *Pseudomonas aeruginosa*. We deleted the gene encoding PAsmr, *smrPA*, from the PA14 genome using *sacB* counterselection to generate an unmarked strain (Figure 6A). We then compared growth of WT PA14 and the PA 14 Δ*smrPA* in Nutrient Broth. There was no significant growth defect in the Δ*smrPA* strain compared to wild type when cultured in NB alone (Figure 6B). We then tested several known and recently discovered PAsmr substrates in PA14 and the Δ*smrPA* strain. Li *et al.* previously demonstrated that deletion of *smrPA* (then known as *EmrE_Pae_*) decreased resistance to ethidium bromide, acriflavine, and several aminoglycoside antibiotics in PAO1, a less virulent strain of *P. aeruginosa* (6). Deletion of *smrPA* did not result in growth differences in the presence of methyl viologen, or acriflavine near their minimum inhibitory concentrations (MICs), consistent with the results of Li *et al* (7). Deletion of *smrPA* from PA14 substantially increased susceptibility to ethidium bromide at the tested concentration (Figure 6D), consistent with the published results (7). To confirm that *smrPA* expression was responsible for this phenotype, we reintroduced *smrPA^+^* in the knockout strain under control of an IPTG inducible promotor, P*_trc_*. In the presence of 1 mM IPTG, resistance to ethidium was restored (Figure 6C). The Δ*smrPA* strain also showed increased susceptibility to chelerythrine chloride (Figure 6E), consistent with the resistance conferred by PAsmr in *E. coli* (Figure 2C). This confirms that chelerythrine chloride is a substrate of PAsmr and the transport activity contributes measurably to resistance to this compound.

**Figure 6.**
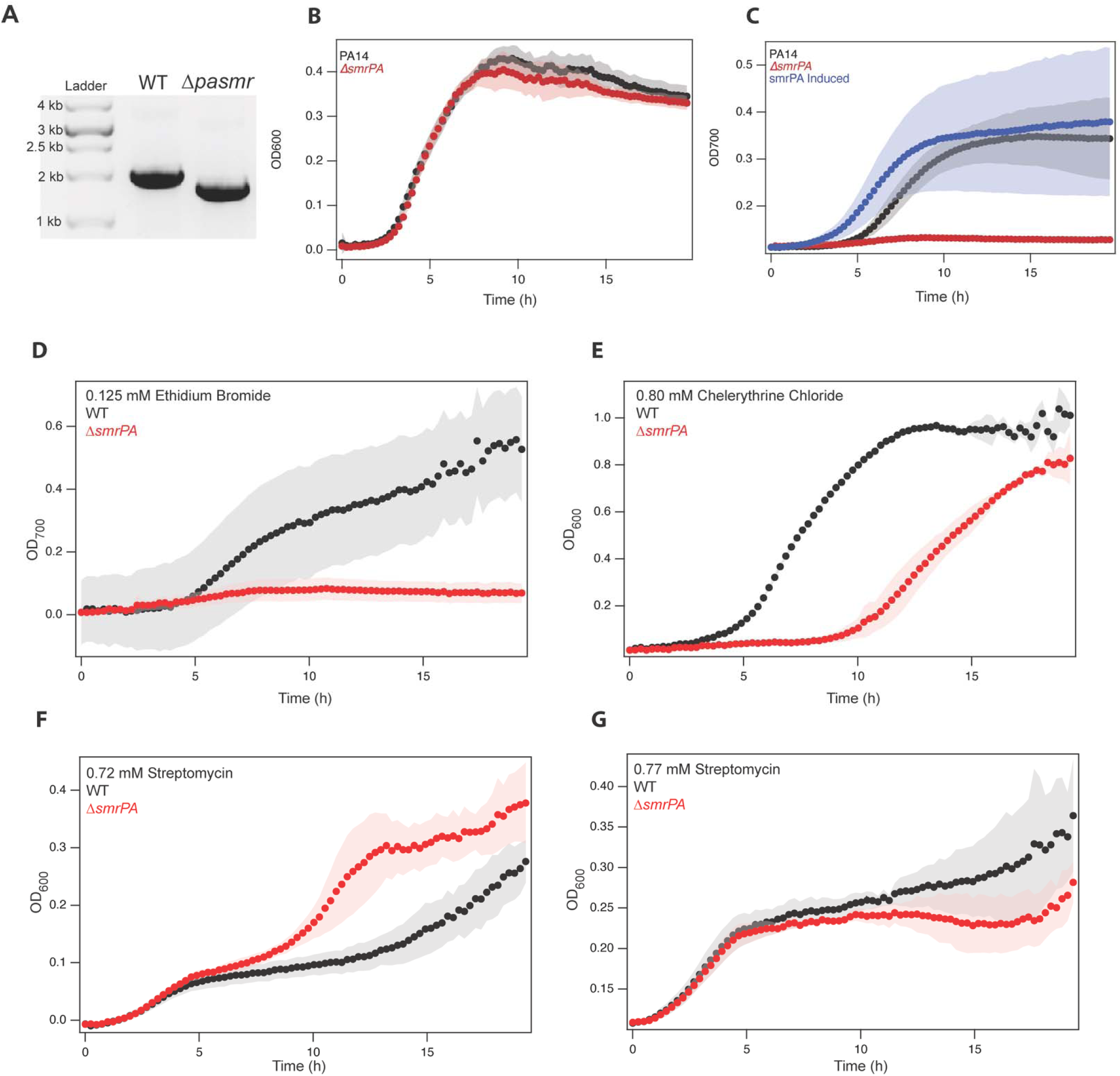
*P. aeruginosa* Δ*smrPA* phenotype is substrate-dependent. A) Confirmation of the *smrPA* deletion in PA14 by PCR with flanking primers. B) Δ*smrPA* does not impact growth in nutrient broth alone. C) Δ*smrPA* with smrPA induced under a different promotor can confer resistance to ethidium bromide. D) Δ*smrPA* is more susceptible to ethidium bromide. E) Δ*smrPA* is more susceptible to chelerythrine chloride. F, G) Δ*smrPA* has a smaller and more varied impact on growth in the presence of streptomycin.

18-crown-6-ether, harmane and streptomycin trigger proton leak through PAsmr that is detrimental to bacterial growth in *E. coli.* However, 18-crown-6 ether was not bactericidal to PA14 or PA14 Δ*smrPA* even at high concentrations (10 mM) and no difference in growth was observed between the two strains (Figure S11F). Upon testing harmane and streptomycin, we observed small and somewhat variable differences between PA14 and PA14 Δ*smrPA* (Figure 6F,G, S11A-D). For each of these substrates, the experiment was performed in triplicate on 3 different days (9 replicates total). For harmane, no significant difference was observed in two out three trials (Figure S11A,B), and a resistance phenotype was observed in the third trial only after a very long lag in growth (Figure S11C). The Δ*smrPA* strain showed increased susceptibility to harmane at 2 mM, but not at lower concentrations (Fig. S11A, B, and C). Streptomycin showed a small resistance phenotype in two out of three trials (Figure 6F, S11D) and no significant phenotype in the third trial (Figure 6G). We also tested tobramycin, a clinically relevant aminoglycoside, for comparison with streptomycin and saw no significant phenotype in PA14 (Figure S11E). The small and less consistent phenotypes observed for harmane and 18-crown-6-ether in *P. aeruginosa*, suggest that substrate-triggered proton leak through PAsmr has less of an impact on this organism. Thus, *P. aeruginosa* responds differently and may be better able to compensate for proton leak than *E. coli*.

## DISCUSSION

Our model of SMR transporter function predicts that these efflux pumps act as double-edge swords for bacteria by conferring resistance to some compounds and susceptibility to others. This investigation serves as a starting point for the further characterization of PAsmr as well as the molecular determinants of substrate profile and phenotype for SMR transporters more broadly. While PAsmr was identified based on sequence homology to EmrE, there are limited data on its function and mechanism. The data presented here confirm that PAsmr transport activity confers resistance to the previously identified substrate, ethidium, and to chelerythrine chloride, which was previously not known to be a substrate of PAsmr. Currently known PAsmr substrates, phenotypes, and transport modes are summarized in Table 2.

**Table 2.** PAsmr Substrates and Phenotypes.

| Substrate/Compound | Interaction |
| --- | --- |
| Tetraphenylphosphonium | Inhibits methyl viologen transport *in vitro* (4) |
| Benzalkonium | Inhibits methyl viologen transport *in vitro* (4); PAsmr inhibition re-sensitizes PAsmr-E. coli (23) |
| Acriflavine | Inhibits methyl viologen transport *in vitro* (4); *smrPA* deletion from PAO1 decreases MIC (7) |
| Ethidium | Inhibits methyl viologen transport *in vitro* (4); Transports/effluxes (23); *smrPA* deletion from PAO1 decreases MIC (7); *smrPA* deletion from PA14 decreases resistance; WT expression in E. coli confers resistance; transported *in vitro* via antiport |
| Methyl viologen | Transported *in vitro* (4), weak resistance phenotype (24) |
| Cetylpyridinium | PAsmr inhibition re-sensitizes PAsmr-E. coli (23), WT expression in E. coli confers resistance (24) |
| Cetyltrimethylammonium | PAsmr inhibition re-sensitizes PAsmr-E. coli (23) |
| Gentamicin | *smrPA* deletion from PAO1 decreases MIC (7) |
| Kanamycin | *smrPA* deletion from PAO1 decreases MIC (7) |
| Neomycin | *smrPA* deletion from PAO1 decreases MIC (7) |
| Nalidixic acid | *smrPA* deletion from PAO1 decreases MIC (7) |
| Chelerythrine | *smrPA* deletion from PA14 decreases resistance; WT expression in E. coli confers resistance; transported *in vitro* via antiport |
| Harmane | WT expression in E. coli confers susceptibility; triggers proton uniport |
| 18-crown-6 ether | WT expression in E. coli confers susceptibility; triggers proton uniport |
| Streptomycin | WT expression in E. coli confers susceptibility; binds |
| Methyltriphenylphosphonium | WT expression in E. coli confers resistance; transported *in vitro* via |
|  | antiport |

Recently, we discovered that EmrE can confer resistance to some substrate and susceptibility to others by performing different types of coupled and uncoupled transport depending on the identity of the substrate (15). Proton-coupled antiport of toxic compounds leads to active efflux and confers resistance to those substrates, the well-known function of the SMR transporter family (Figure 3A).

Harmane triggers uncoupled proton uniport through EmrE, dissipating the proton gradient and negatively impacting ATP levels and growth in *E. coli*. The solid-supported membrane electrophysiology data in Figure 3 show that both harmane and 18-crown-6-ether trigger proton leak through PAsmr, the first direct confirmation that drug-gated proton uniport occurs in an SMR transporter other than EmrE. When PAsmr is heterologously expressed in *E. coli*, this transport activity is similarly detrimental to bacterial growth and ATP levels – *E. coli* expressing non-functional transporter have higher ATP levels than cells expressing functional transporter, and functional transporter also impairs growth.

The discovery that the aminoglycoside antibiotic streptomycin is a substrate of PAsmr supports the need for greater understanding of SMR transporters in clinically relevant organism, and raises further questions about the impact of proton leak. 18-crown-6 ether and harmane trigger proton leak, but the mechanism by which PAsmr induces susceptibility to streptomycin in *E. coli* remains unconfirmed, although streptomycin does bind to PAsmr (Figure 4). While 18-crown-6 ether and harmane trigger a consistent decrease in ATP production, *E. coli* expressing PAsmr were able to eventually recover growth in the presence of streptomycin, leading to *increased* ATP production compared to control by the time they reached stationary phase (Figure 2G). This may be explained by PMF-mediated uptake of aminoglycosides or the previous observation that streptomycin itself causes time dependent changes in ATP levels due to its well-known function suppressing protein synthesis (35). Thus, an alternative possibility is that streptomycin-triggered proton leak through PAsmr reduces ATP level initially and then this is overcome by the reduced use of ATP in protein production upon streptomycin targeting of the ribosome.

When moving to the native organism, *P. aeruginosa*, we observed significant differences in the consistency of PAsmr to confer resistance or susceptibility across different organisms. In the case of proton-coupled antiport mediated resistance to toxic compounds, the function of PAsmr was consistent in *E. coli* and *P. aeruginosa.* While Li *et al.* demonstrated that *smrPA* deletion results in a reduced minimum inhibitory concentration to some substrates in PAO1, only traditional transport substrates and aminoglycosides were tested, and full growth curves were not examined (7). We demonstrated that deletion of *smrPA* from PA14, a more virulent model strain, resulted in loss of resistance to ethidium bromide compared to WT, which is surprising given the numerous additional efflux mechanisms present in this strain (39, 40).

However, the difference in growth between WT PA14 and PA14 Δ*smrPA* with the three tested susceptibility substrates was more subtle and inconsistent. This highlights important nuances in how proton leak impacts bacterial growth in different organisms and prompts further investigation about the greater ability of *P. aeruginosa* to compensate for or prevent loss of ΔpH due to transporter-mediated proton leak compared to *E. coli*. As PAsmr did confer resistance to ethidium and chelerythrine, the more subtle phenotype with susceptibility substrates is not due to lack of PAsmr activity. Using PAsmr to trigger proton leak provides a pathway to disrupt ΔpH specifically without more general membrane disruption, in constrast to small molecule PMF disruptors that act via the membrane, such as nigericin or CCCP. Thus, triggering proton leak through PAsmr provides a new route to probe the mechanisms by which *P. aeruginosa* responds to specific PMF disruption.

Although the growth phenotypes were subtle and somewhat variable, the ability to induce proton leak through PAsmr may still provide a useful route to develop compounds that can synergize with other drugs that are dependent on the PMF or impacted by the metabolic response of *P. aeruginosa* to ΔpH dissipation. Importantly, targeting PAsmr in this manner, by triggering an alternative transport behavior rather than inhibiting transport, does not require PAsmr to be the primary resistance mechanism, but simply be present and expressed in a particular *P. aeruginosa* strain or isolate. Given that SMR transporter orthologs are generally conserved in *P. aeruginosa* isolates (nearly 300 *P. aeruginosa* strains in the pseudomonas.com database contain SMR transporter orthologs [Winsor: 26578582]), the demonstration that PAsmr is capable of substrate-induced proton uniport makes it a relevant target. Regulation of efflux pumps is a key resistance mechanism in P. *aeruginosa*, and understanding the expression of transporters, including SMR transporters is also an important future direction. Ultimately, understanding the organismal response to diverse transport behavior in complex resistance networks will aid future antibiotic development efforts.

## SUPPORTING INFORMATION (ATTACHED)

Supplemental Methods Fig. S1-S11, Table S1

Data repository: https://data.mendeley.com/preview/nng7769fbj (until published)

## ACKNOWLEDGEMENTS

The authors wish to thank Grant A. Hussey for aiding in the initial plasmid design of the pWB vector and Kylie M. Hibbs for preliminary growth curve work. Thank you to Andrew Buller and members of the Henzler-Wildman and Peters lab for thoughtful comments. Mass spectrometry was carried out by the University of Arizona Analytical & Biological Mass Spectrometry Facility (RRID:SCR_023370). Research reported in this publication was supported by the National Institutes of Health Institute for General Medical Sciences under award numbers R01GM095839 and R35GM141748, with additional support for AKW from the Institute of Allergy and Infectious Diseases under award number F31AI169825. WJH and AKW were also supported by the Biotechnology Training Program (NIH 5T32GM135066). This study made use of the National Magnetic Resonance Facility at Madison, which is supported by NIH grant R24GM141526. The content is solely the responsibility of the authors and does not necessarily represent the official views of the NIH.

## AUTHOR CONTRIBUTIONS

Conceptualization-KAHW, JMP, AKW; Methodology-KAHW, JMP, WJH, AKW; Investigation-AKW, WJH, SPD, MSM; Data Curation-AKW, WJH, SPD, MSM; Writing-Original Draft-AKW, MSM; Writing-Review & Editing-AKW, KAHW, JMP, WJH; Visualization-AKW, MSM; Supervision-KAHW, JMP; Funding acquisition-KAHW, AKW

