## Supplemental Material for "A Small Multidrug Resistance Transporter in *Pseudomonas aeruginosa* Confers Substrate-Specific Resistance or Susceptibility"

**This file includes:**

Supplementary Fig. 1-11

Supplementary Table 1

**Additional data:**

doi:10.17632/nng7769fbj.1 (<https://data.mendeley.com/preview/nng7769fbj> until published)

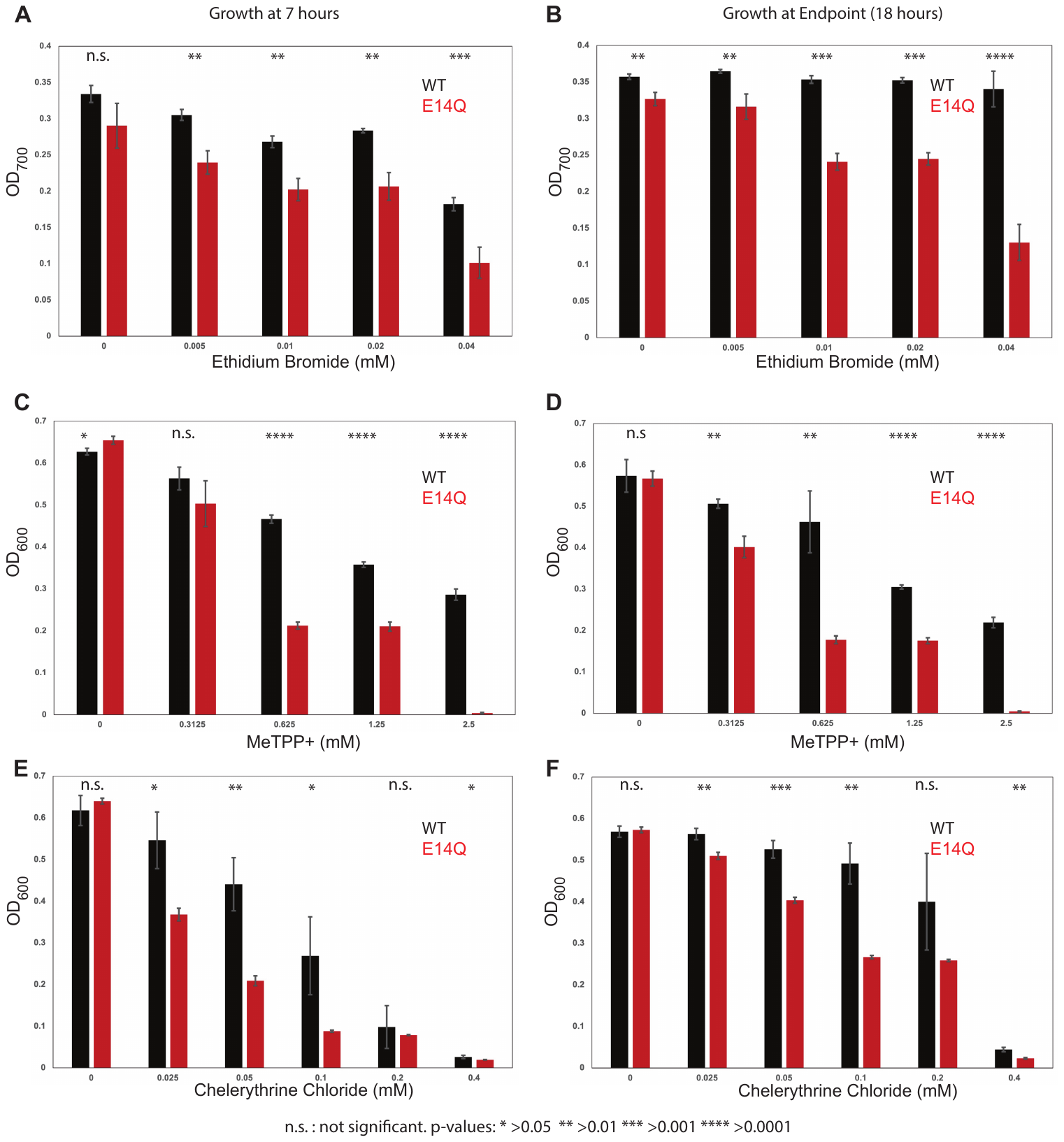
**Figure S1. Concentration dependence of resistance substrates in *E. coli*.** At 7 hours (A, C, E) and 18 hours (B, D, F), *E. coli* expressing WT PAsmr mostly show increased growth compared to the transport-dead control mutant E14Q. This effect is concentration-dependent, typically with greater significance at higher concentrations. Error bars represent standard deviation of replicates.

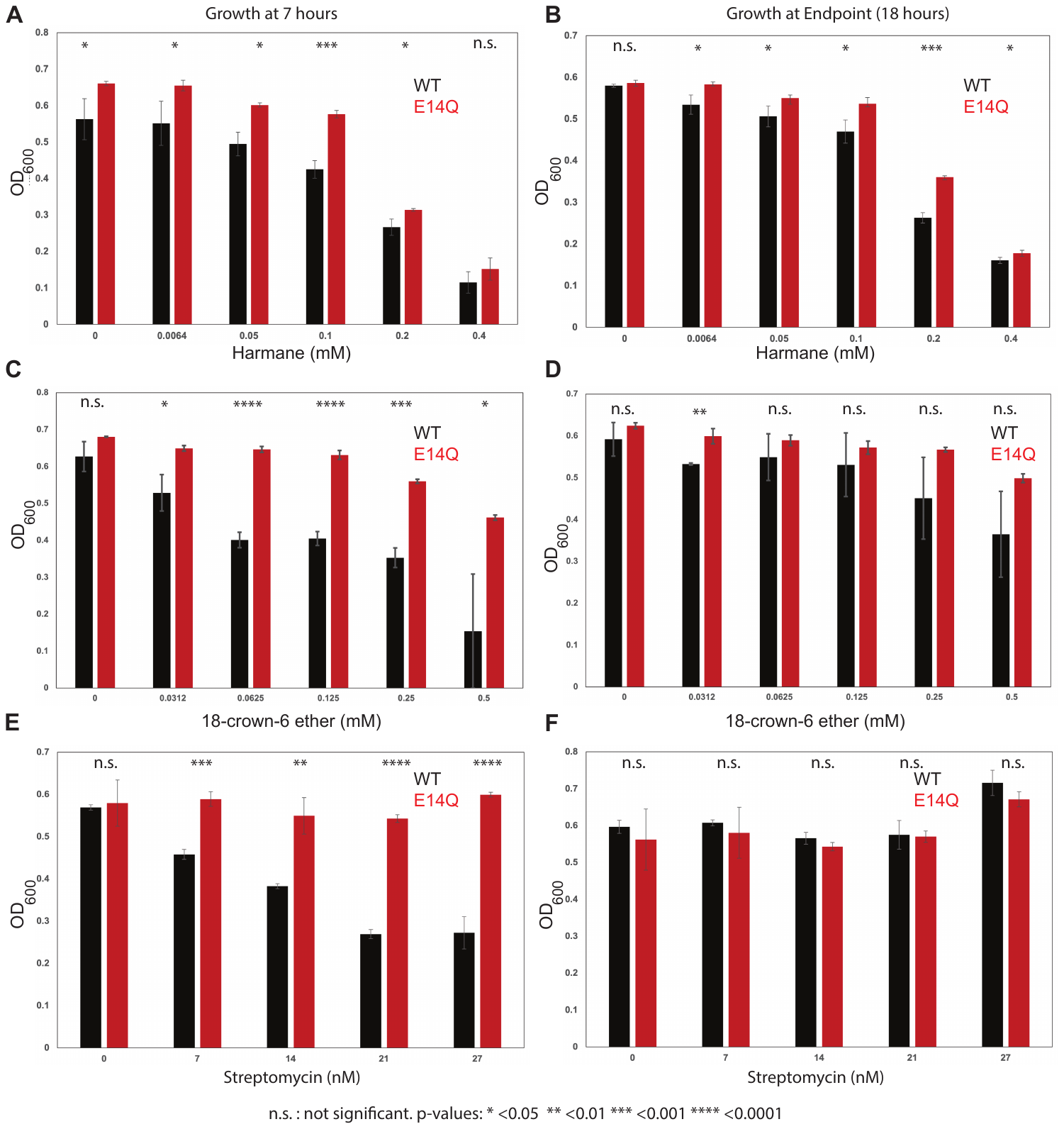
**Figure S2. Concentration dependence of susceptibility substrates in *E. coli.*** At 7 hours (A, C, E) and 18 hours (B, D, F), *E. coli* expressing WT PAsmr mostly show decreased growth compared to the transport-dead control mutant E14Q. This effect is concentration-dependent, typically with greater significance at higher concentrations. For streptomycin (E, F), the phenotype is pronounced at 7 hours but growth of WT-PAsmr cells has recovered by the endpoint. Error bars represent standard deviation of replicates.

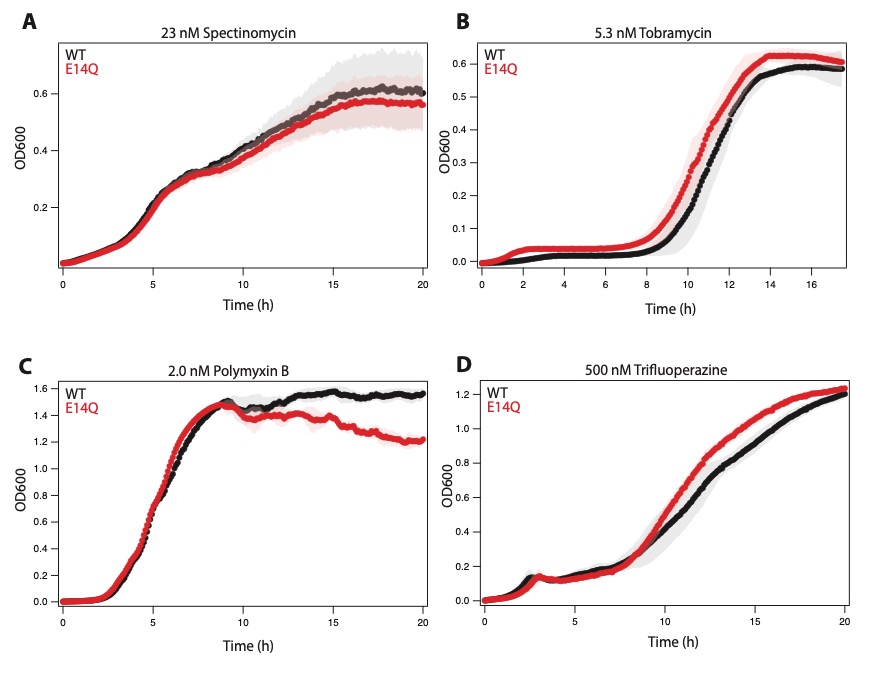
**Figure S3. Compounds to which PAsmr confers no growth phenotype.** Several substrates from the screen in (24) were tested but functional PAsmr did not confer a difference in growth in *E. coli* compared to control*.* Tobramycin (B) was also tested due to similarity to streptomycin. All trials were conducted with n = 3 and data at the first sub-lethal concentration are shown unless otherwise noted.

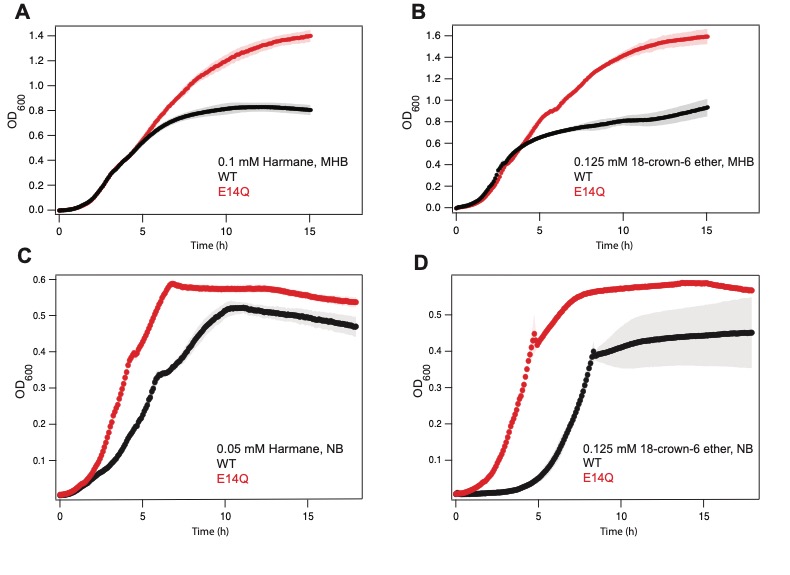

**Figure S4. Impact of media ionic strength on susceptibility phenotype.** In Muller-Hinton Broth, a high ionic strength medium, susceptibility phenotypes of harmane (A) and 18-crown-6 ether (B) do not appear until around 6 hours. These experiments were published in (24). In Nutrient Broth (see also Figure 2), harmane (C) and 18-crown-6 ether (D) phenotypes are more quickly apparent, although result in a lesser difference in carrying capacity at endpoint**.**

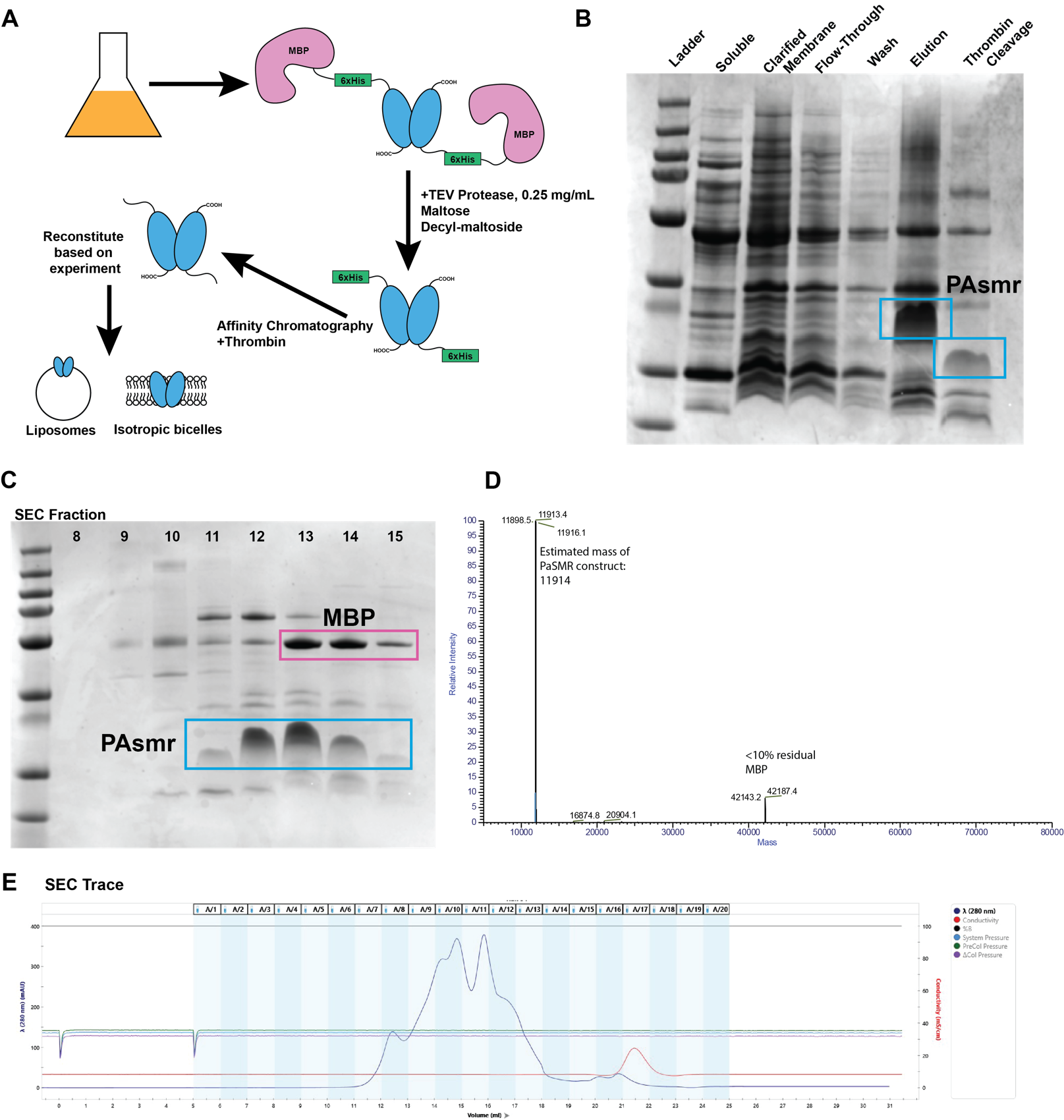

**Figure S5. Expression and Purification of PAsmr for biophysical study.** A) Purification scheme of PAsmr expressed as a fusion with Maltose Binding Protein (MBP) and an N-terminal 6xHis tag. B) Purification protocol results in elution of PAsmr off NiNTA column and successful cleavage of 6xHis tag. C) Size-exclusion chromatography separates PAsmr from residual MBP. D) Intact Mass Analysis Mass Spectrometry showing mass consistent with PAsmr. E) Size-exclusion chromatogram.

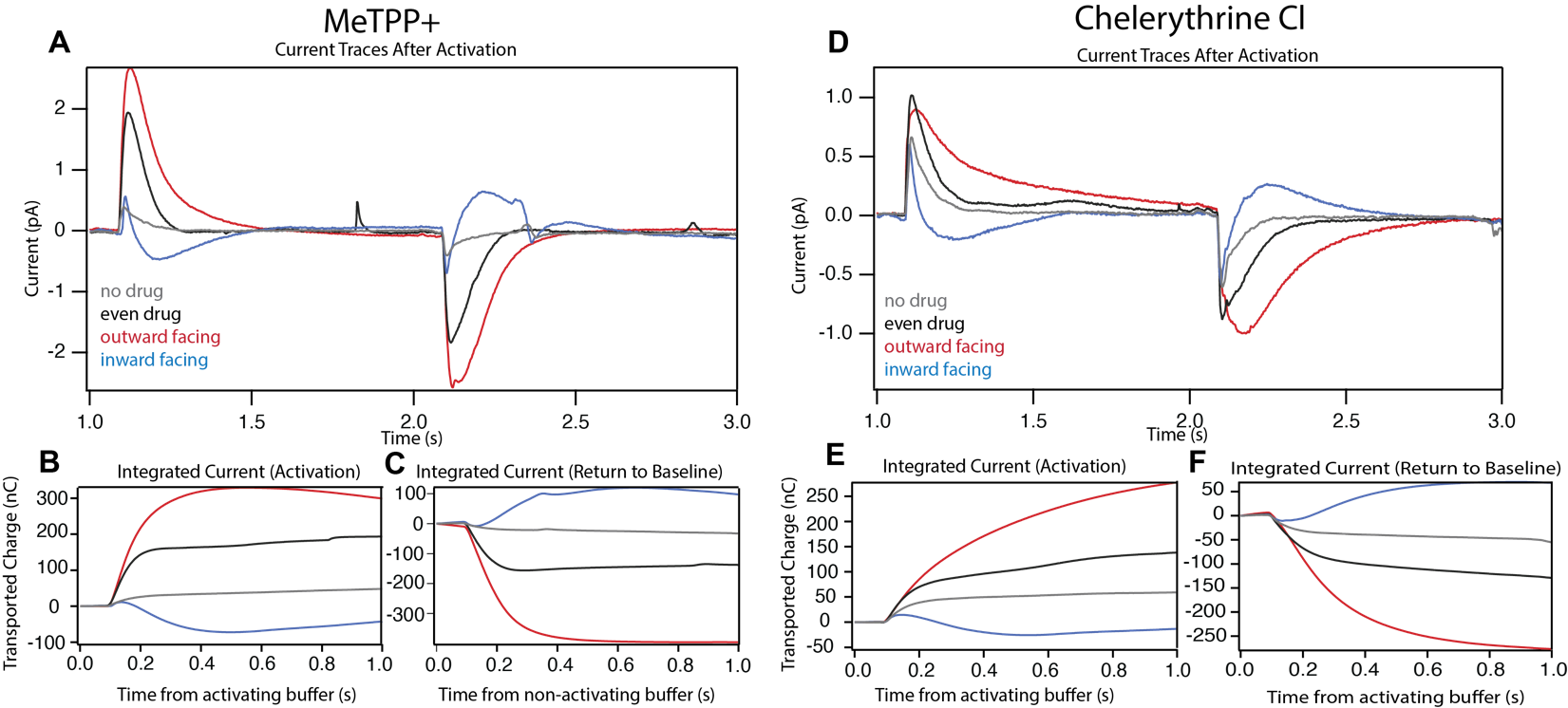

**Figure S6. SSME current traces of resistance substrates.** Raw (A, D) and Integrated (B, C) average traces for each gradient condition for MeTPP+ and Chelerythrine Cl-.Activating buffer is introduced at 1 second and non-activating buffer is reintroduced at 2 seconds.

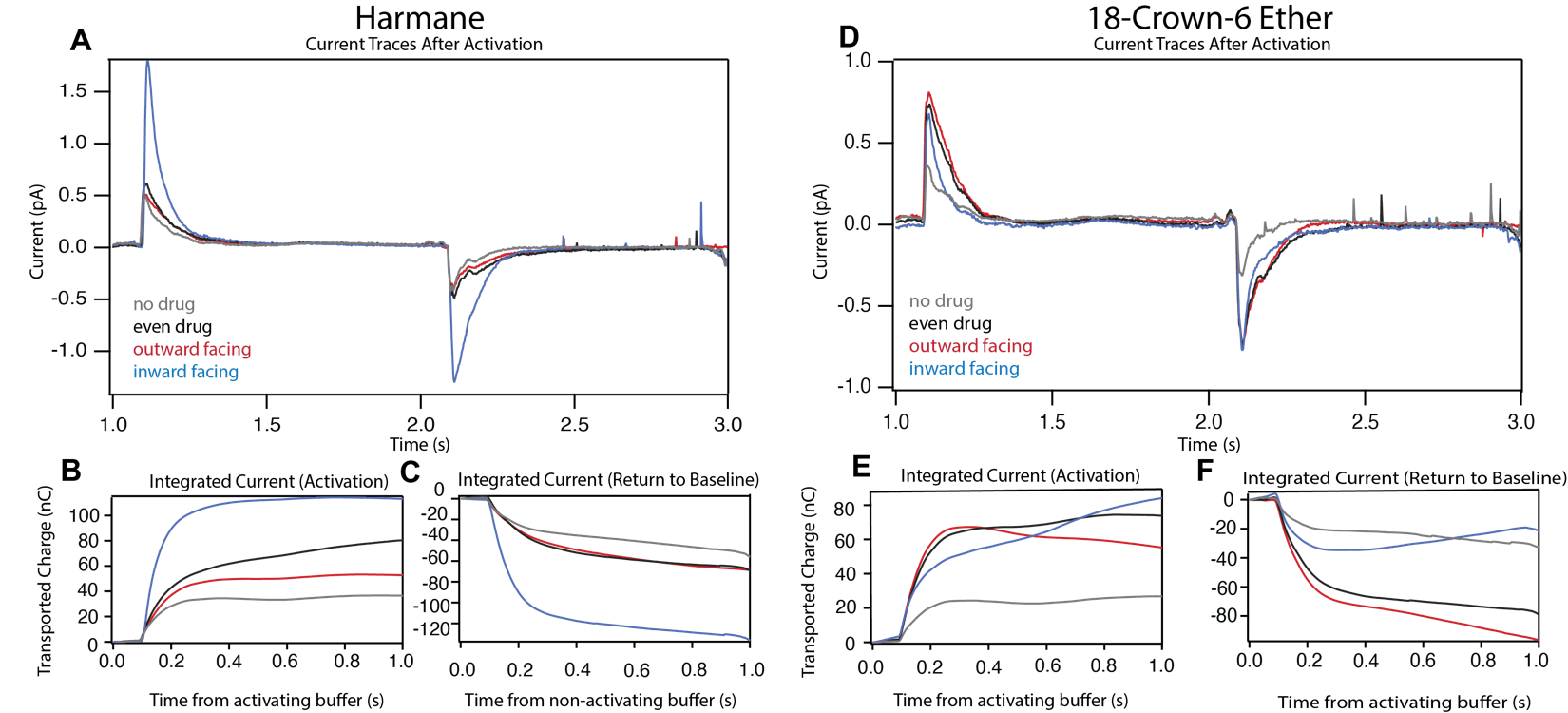

**Figure S7. SSME current traces of susceptibility substrates.** Raw (A, D) and Integrated (B, C) average traces for each gradient condition for Harmane and 18-crown-6 ether. Activating buffer is introduced at 1 second and non-activating buffer is reintroduced at 2 seconds.

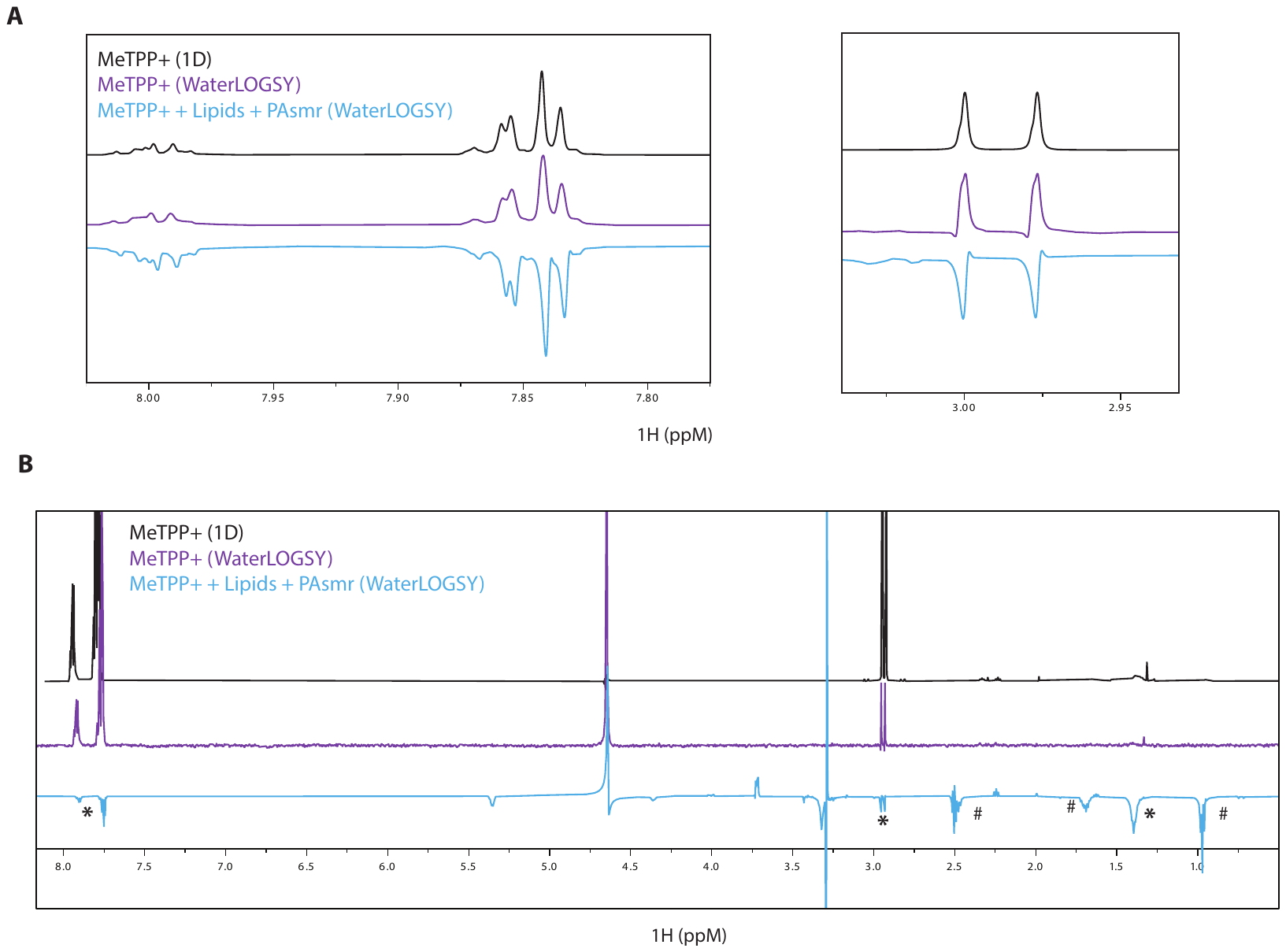

**Figure S8. 1D Ligand-detected NMR shows binding of MeTPP^+^.** Addition of PAsmr to MeTPP^+^ results in a negative WaterLOGSY signal for the MeTPP^+^ resonances (marked with *), while the signal is positive when only lipids are present. This confirms that MeTPP^+^ interacts specifically with PAsmr and that WaterLOGSY analysis of PAsmr is effective. Interestingly, the lipid signals are also negative in the WaterLOGSY spectra (marked with #) when PAsmr is present, suggesting lipid binding to PAsmr. Key regions (A) and full spectra (B) are shown and full spectra can also be found in data repository.

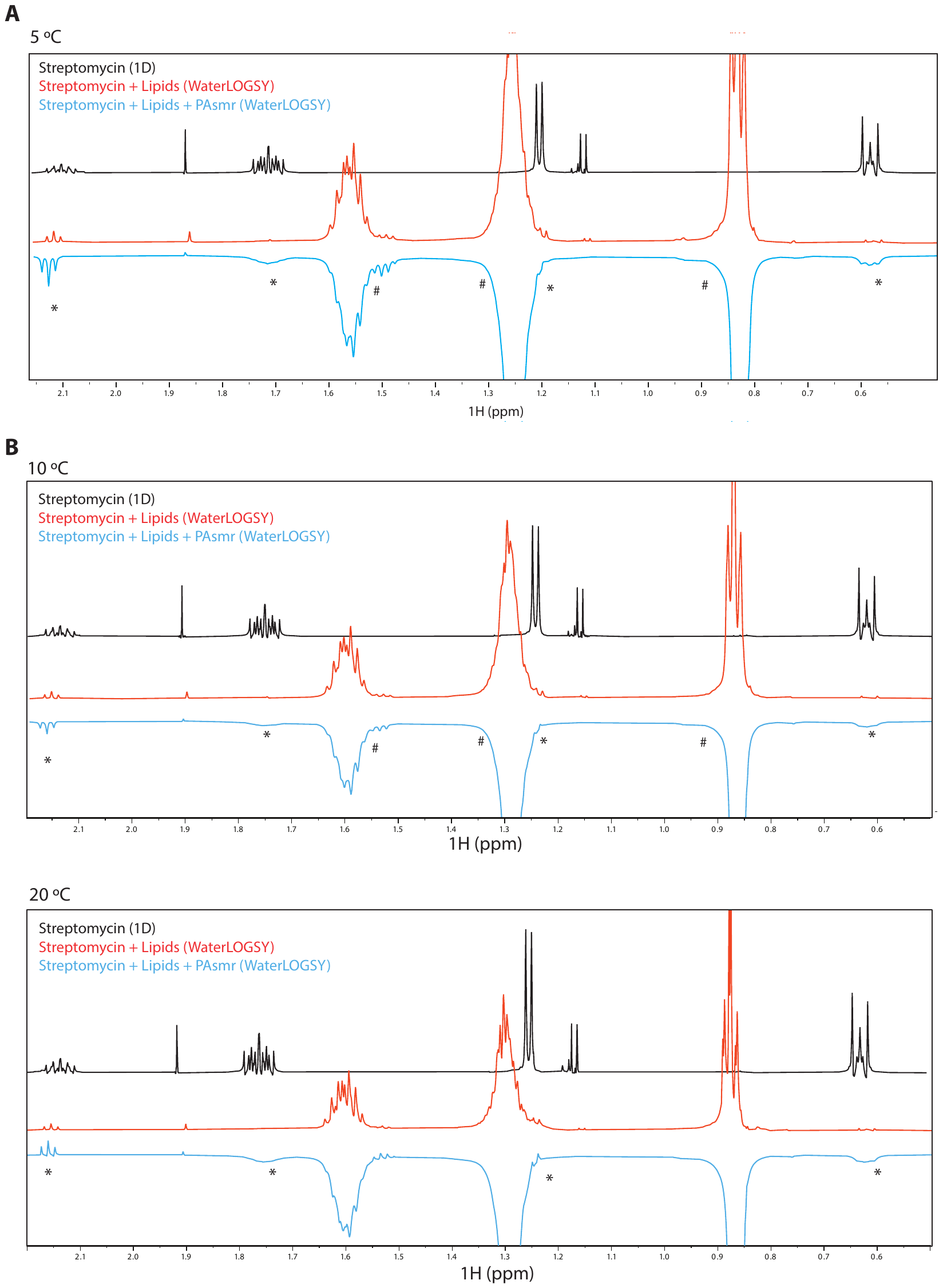
**Figure S9. WaterLOGSY binding effect at 5 and 10 degrees Celsius.**

Addition of PAsmr to streptomycin results in negative signal at characteristic streptomycin chemical shifts (*). Interestingly, the lipid signals are negative in the WaterLOGSY spectra when PAsmr is present (#), suggesting lipid binding to PAsmr. Representative regions of spectra are shown and full spectra can be found in data repository.

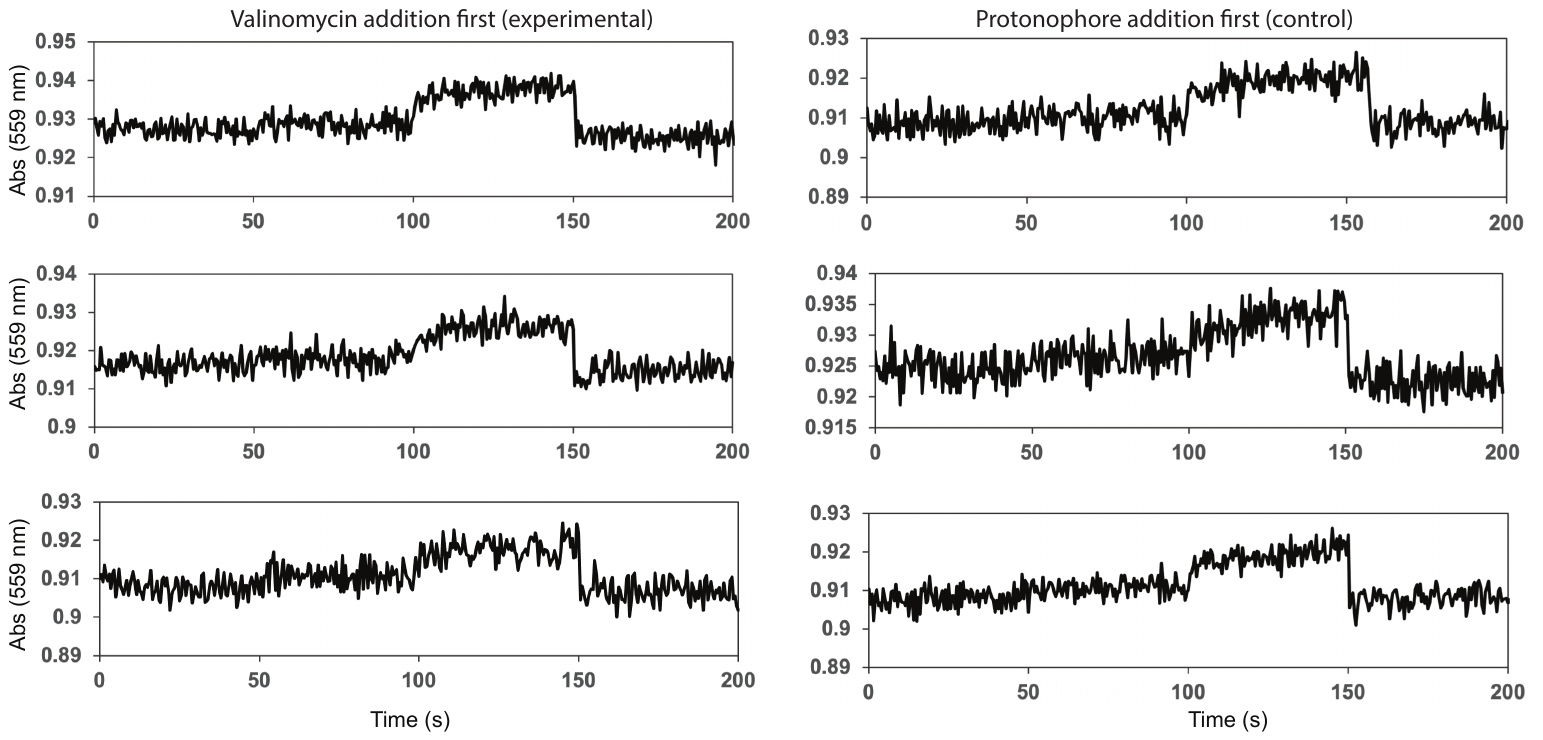

**Figure S10. All proton leak traces.** Top traces are included in Figure 5. Additional replicates are second and third row. For experimental set, valinomycin was added around 50 s, protonophore around 100 s, and HCl around 150 s. For control set, protonophore was added around 50 s, valinomycin around 100 s, and HCl around 150 s.

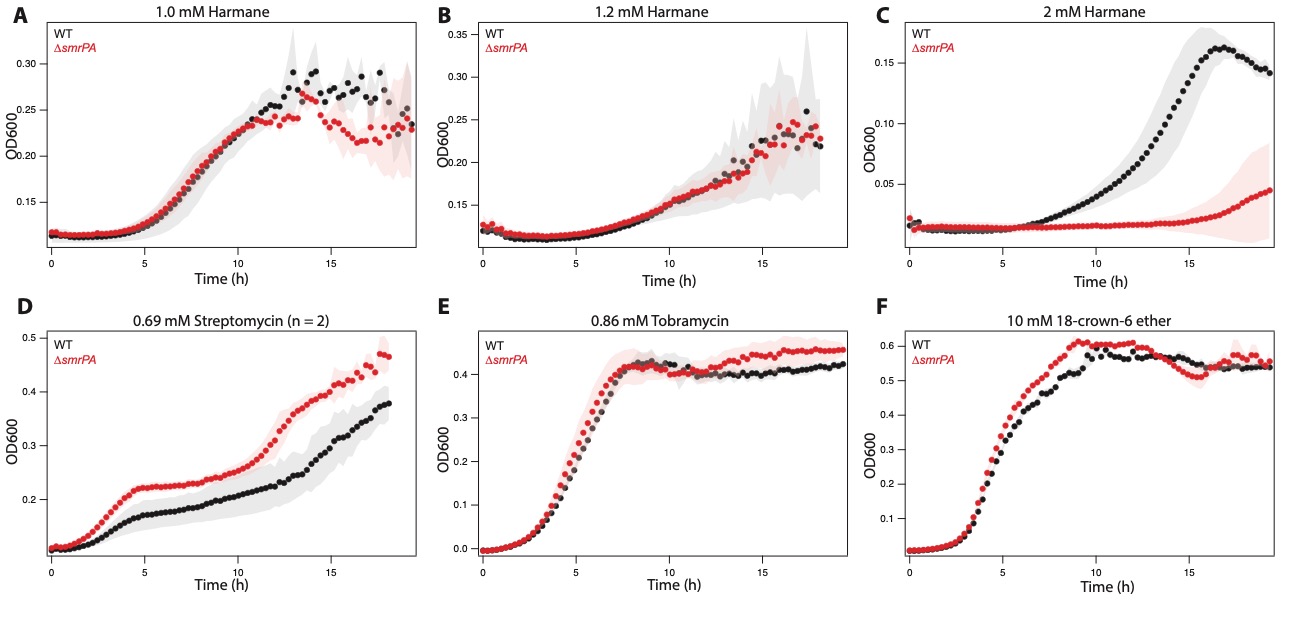
**Figure S11. Additional *∆smrPA* growth curves.** A-C) Growth in harmane at high concentrations typically did not differ between PA14 and the knock-out, but occasionally WT grew better compared to the knock-out. D) Additional trial of streptomycin, with only 2 biological replicates. E) *smrPA* deletion did not result in a phenotypic difference in the presence of tobramycin. F) 18-crown-6 ether did not inhibit growth of PA14 at even very high concentrations. All trials were conducted with n = 3 and data at the first sub-lethal concentration are shown unless otherwise noted.

**Table S1. Concentrations used in SSME**

| **Compound** | **High Concentration (µM)** | **Low Concentration (µM)** |
| --- | --- | --- |
| MeTPP+ | 1 | 0.032 |
| Chelerythrine | 4 | 0.63 |
| Harmane | 4 | 0.124 |
| 18-crown-6 ether | 320 | 10 |
